# BNIP3L/NIX-mediated mitophagy limits Alzheimer’s disease pathology and preserves neuronal function

**DOI:** 10.64898/2026.10.18.759942

**Authors:** Yuan Li, Christina ploumi, Christiane Volbracht, Weihua Tian, Nefeli Kalogeraki, Katerina Kitopoulou, Claudia Carvalho Navarro, Zhiquan Li, Nicolas Cherbuin, Leticia Camargo Tavares, Zilu Ye, Aleksander J. Norwak, Satyan Chintawar, M Zameel Cader, Laura Berliocchi, Steffen Goletz, Vilhelm A. Bohr, Kristine Freude, Konstantinos Palikaras, Lene Juel Rasmussen

**Author notes:** These authors contributed equally to this work (Yuan Li, Christina ploumi).

## Abstract

Defective mitophagy is emerging as a key contributor to Alzheimer’s disease (AD), but the neuronal pathways governing mitochondrial clearance and their contribution to disease progression remain poorly understood. Here, we identify BNIP3L/NIX-dependent mitophagy as a conserved neuronal quality control pathway that maintains mitochondrial fitness and constrains AD-associated proteotoxicity. Loss of BNIP3L/NIX or its *C. elegans* orthologue DCT-1 impaired basal and stressed-induced mitophagy, disrupted mitochondrial homeostasis, enhanced Tau and Aβ pathology and compromised neuronal network, synaptic and behavioral function across human neuronal and *in vivo* models. Conversely, restoring BNIP3L/NIX and DCT-1 activity improved neuronal function under proteotoxic stress *in vivo*, reduced disease-associated Tau phosphorylation in human neurons and markedly suppressed seeded Tau aggregation in primary mouse neurons. Together, these findings identify BNIP3L/NIX-mediated mitophagy as a mechanistic link between mitochondrial quality control, proteostasis and neuronal function, and establish its restoration as a potential strategy for counteracting AD-associated pathology.

## Introduction

Alzheimer’s disease (AD) is characterized by extracellular amyloid-β (Aβ) deposition, intracellular accumulation of hyperphosphorylated Tau, progressive synaptic dysfunction and cognitive decline^1, 2^. Mitochondrial dysfunction emerges early in AD and is increasingly recognized as an important contributor to disease pathogenesis^3–9^. Aβ and Tau disrupt mitochondrial homeostasis, while mitochondrial dysfunction, in turn, can promote pathological protein accumulation^10–12^, suggesting a reciprocal relationship between proteotoxicity and impaired mitochondrial quality control. Defining the mechanisms that preserve a functional mitochondrial pool in neurons are therefore likely to reveal how mitochondrial dysfunction contributes to AD-associated neuronal impairment. Mitophagy, the selective delivery of mitochondria to lysosomes for degradation, is essential for mitochondrial quality control and homeostasis^13, 14^. Mitophagy is impaired during aging and AD, whereas genetic or pharmacological enhancement of mitochondrial clearance can attenuate Aβ and Tau pathology and improve cognitive function in experimental AD models^13–15^. Much of the mechanistic understanding of mitophagy has centered on the ubiquitin-dependent PINK1–PARKIN pathway. However, studies using mitophagy reporter models have shown that basal mitophagy persists in the absence of PINK1 or PARKIN in several tissues and neuronal populations *in vivo*, indicating that constitutive mitochondrial turnover can occur independently of this pathway^16–18^. Receptor-mediated mitophagy provides an alternative mechanism by which mitochondrial outer membrane proteins directly engage the autophagic machinery^19, 20^. Yet, which mitophagy receptors sustain mitochondrial turnover in neurons, and whether their dysfunction contributes directly to AD-associated proteinopathy and neuronal dysfunction remain poorly understood.

BNIP3L (also known as NIX; hereafter BNIP3L) is a mitochondrial outer membrane mitophagy receptor that promotes selective mitochondrial clearance through direct engagement of the autophagic machinery. Beyond its established role in erythrocyte maturation^21^, BNIP3L-dependent mitophagy contributes to mitochondrial remodeling during neuronal differentiation^20, 22^. In *Caenorhabditis elegans (C. elegans)*, its homologue DCT-1 coordinates mitophagy and mitochondrial biogenesis^23^, while BNIP3-family-mediated mitophagy promotes mitochondrial homeostasis and longevity in *Drosophila melanogaster*^24^. Furthermore, BNIP3L-mediated mitophagy has also been implicated in neuronal stress responses and synaptic maintenance^25^. These observations suggest that BNIP3L can provide an important receptor-mediated mechanism for neuronal mitochondrial quality control. However, whether BNIP3L sustains basal and stress-induced mitophagy in neurons, and whether failure of this pathway contributes to AD-associated proteotoxicity and neuronal dysfunction, remain unknown.

Here, we identify BNIP3L-mediated mitophagy as a conserved neuronal quality control pathway linking mitochondrial homeostasis to AD-associated proteinopathy and neuronal function. Using complementary human induced pluripotent stem cell (iPSC)-derived neurons, primary mouse neurons, and *C. elegans* models, we show that disruption of BNIP3L/DCT-1 impairs neuronal mitophagy and mitochondrial fitness, promotes Tau and Αβ pathology and compromises neuronal network, synaptic and behavioral function. Conversely, restoration of BNIP3L/DCT-1 activity improves neuronal function and suppresses pathological Tau accumulation across experimental systems. Together, our findings establish receptor-mediated mitophagy as an important determinant of neuronal resilience and identify BNIP3L/DCT-1 as a conserved pathway coupling mitochondrial quality control to proteostasis and neuronal function in AD.

## Results

### BNIP3L is induced during neuronal differentiation, and its adaptive response is impaired in AD neurons

To determine which mitophagy pathways are preferentially activated during neuronal differentiation, we differentiated human induced pluripotent stem cells (iPSCs) into cortical neurons (Fig. S1a-e). BNIP3L expression increased markedly during differentiation, whereas PINK1, MUL1 and the mitophagy receptor AMBRA1 showed comparatively modest changes (Fig. 1a, b). These findings identify BNIP3L as a mitophagy regulator preferentially induced during neuronal maturation.

**Figure 1.**
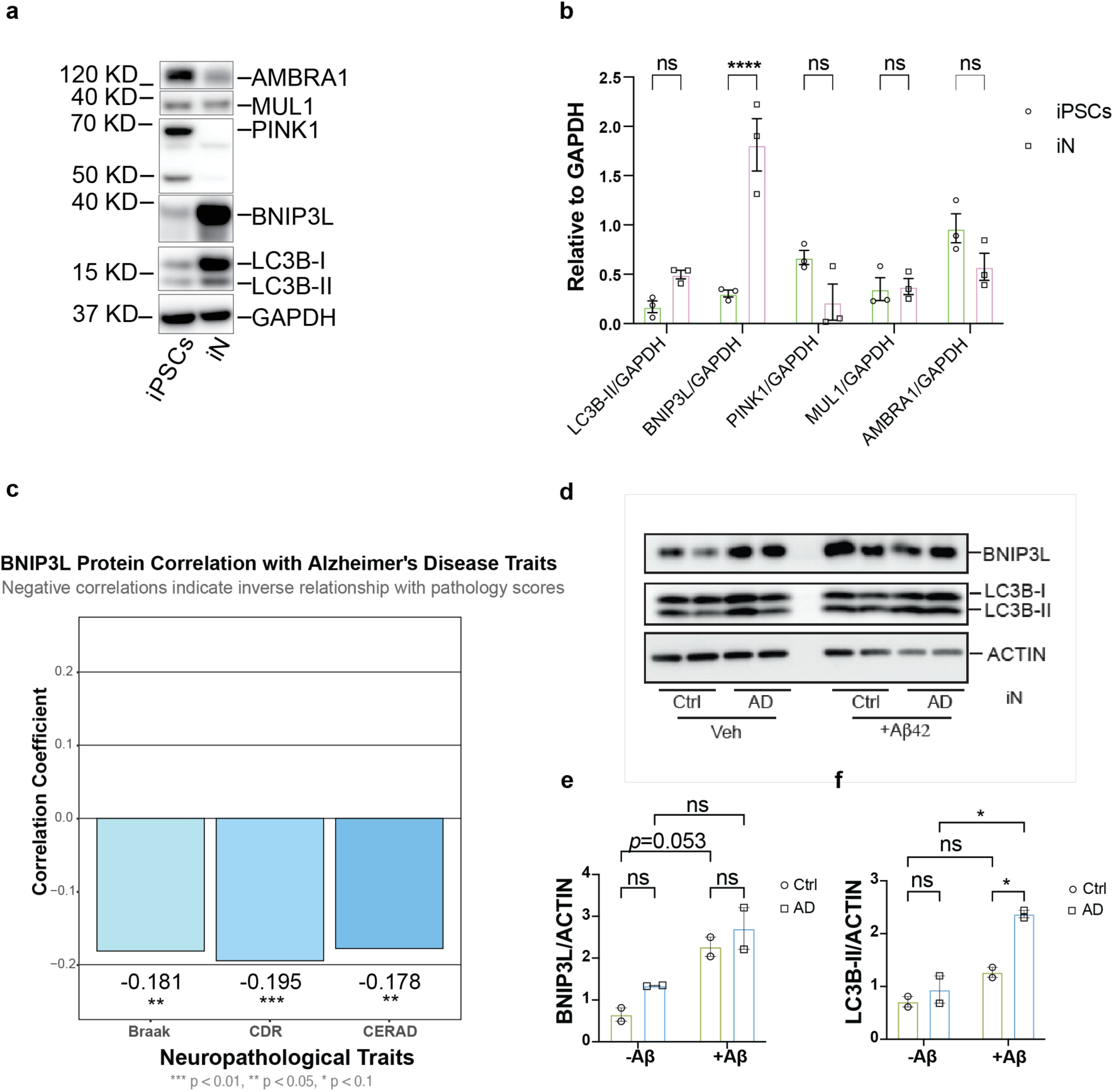
BNIP3L is induced during neuron differentiation and correlates with AD traits. **a.** Representative image of Western blot data showing expression of mitophagy-related proteins including AMBRA1, MUL1, PINK1, BNIP3L and LC3B’s expression in iPSCs and iPSCs-derived neurons (iN). **b.** Quantification of the Western blot data in Figure 1a. Data was obtained from three batches of independent experiments and presented as mean ± SEM. Statistical significance was determined using two-way ANOVA Šídák’s multiple comparisons test. **** p < 0.0001, ns, no significant difference. **c.** In the dataset published by Wang *et al.* (*Cell*, 2025), approximately 5,000 Alzheimer’s disease (AD)-associated proteins, termed proteins correlated with traits (PCTs), were identified based on their associations with three clinical and neuropathological measures: Braak stage, which reflects the progression of neurofibrillary tangle pathology; Clinical Dementia Rating (CDR), which assesses dementia severity and cognitive impairment; and the Consortium to Establish a Registry for Alzheimer’s Disease (CERAD) score, which evaluates neuritic plaque burden. BNIP3L was identified as a PCT whose protein abundance was negatively correlated with AD progression. The figure was generated in RStudio using data from page 2 of Table S2 reported by Wang *et al.* (*Cell*, 2025). **d.** Representative Western blot images showing BNIP3L and LC3B’s expression in healthy controls and sporadic Alzheimer’s disease (sAD) patient iPSC-derived neurons treated with vehicle (Veh) or oligomeric Aβ42 (Aβ42; 2.5 μM, 24 h) treatment. **e and f**: Quantification of the Western blot data. Data are presented as mean ± SEM. Statistical significance was assessed using two-way ANOVA followed by Tukey’s multiple-comparisons test. *P* < 0.05; ns, not significant.

We next asked whether BNIP3L is associated with AD pathology in the human brain. Previous transcriptomic analyses reported reduced BNIP3L expression in sporadic AD (sAD) brain tissue^26^. Therefore, we interrogated the proteomic dataset of Wang et al^27^, which relates brain protein abundance to neuropathological and clinical measures of AD. BNIP3L protein abundance was inversely associated with Braak stage, Clinical Dementia Rating (CDR) and CERAD score (Fig. 1c), linking reduced BNIP3L abundance to greater neuropathological burden and clinical disease severity.

We next examined whether the BNIP3L response to AD-associated stress is altered in patient-derived neurons. BNIP3L and LC3B-II protein levels did not differ significantly between age- and sex-matched control and sAD iPSC-derived neurons under basal conditions (Fig. 1d–f). However, oligomeric Aβ42 induced robust BNIP3L upregulation in control neurons, whereas this response was markedly attenuated in sAD neurons (Fig. 1d–f). Thus, although basal BNIP3L abundance was preserved, sAD neurons showed an impaired ability to engage BNIP3L in response to Aβ-associated stress. Together, these findings show that BNIP3L is engaged during neuronal differentiation and suggest that the capacity to recruit this pathway in response to proteotoxic stress is compromised in AD.

### BNIP3L sustains basal and stress-induced neuronal mitophagy and mitochondrial homeostasis

To determine whether BNIP3L is required for neuronal mitophagy, CRISPR–Cas9-generated *BNIP3L* knockout (KO) iPSCs and isogenic controls (Fig. S2a) were differentiated into neurons. BNIP3L was subsequently re-expressed in KO neurons to examine whether the resulting phenotypes were specifically attributable to BNIP3L loss (Fig. S2b).

We first assessed basal mitochondrial turnover following lysosomal blockade with chloroquine (CQ)^28^. Analysis of LC3B-II together with the mitochondrial proteins MTCO2 and TOMM20 revealed reduced mitophagy flux^29^ in *BNIP3L* KO neurons (Fig. S2c). Consistently, imaging-based analysis showed reduced mitochondria-lysosome colocalization in *BNIP3L* KO neurons compared with WT controls (Fig. 2a). Re-expression of BNIP3L restored mitochondria–lysosome colocalization (Fig. 2a) and mitophagy flux (Fig. S2d), supporting a specific requirement for BNIP3L in basal neuronal mitophagy.

**Figure 2.**
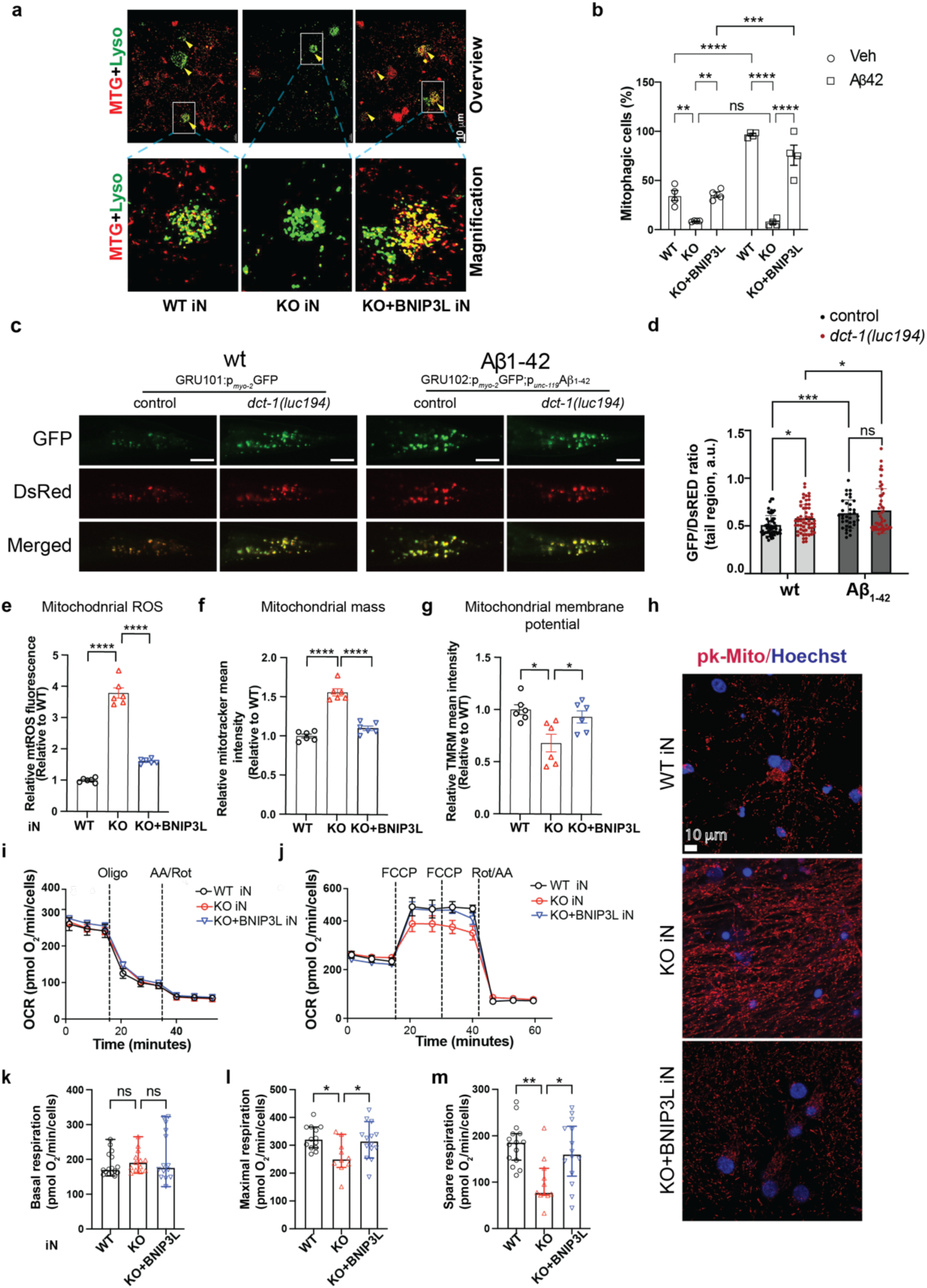
BNIP3L deficiency impairs mitophagy and disrupts mitochondrial homeostasis. **a.** Representative confocal images showing the co-localization of the mitophagy dye (red) and lysotracker dye (green) in WT, KO, and KO+BNIP3L iN. Yellow arrowhead indicates the colocalization of mitophagy dye and lysosome dye. Scale bar, 10 mm. **b.** Percentage of mitophagic cells measured by flow cytometry in mt-Keima–integrated WT, KO, and KO+BNIP3L iN under basal conditions or following oligomeric Aβ42 treatment. Data are presented as mean ± SEM. Statistical significance was determined using two-way ANOVA followed by Šidák’s multiple-comparisons test. ** P < 0.01, ***P < 0.001, and ****P < 0.0001; ns, not significant. **c.** Representative fluorescence images of 1-day-old transgenic *C. elegans* expressing the pan-neuronal mitoRosella reporter (p*_unc-119_*::TOMM-20::Rosella) in wild-type (GRU101) and Aβ-expressing (GRU102) backgrounds under control or *dct-1(luc194)* mutant conditions. Scale bar, 25 μm. **d.** Quantification of neuronal mitophagy based on the GFP/DsRed fluorescence ratio of the mitoRosella reporter measured in the tail region of the nematodes. Lower GFP/DsRed ratios indicate increased mitophagy (data presented as mean ± SD; ns p > 0.05, *p < 0.05, ***p < 0.001; one-way ANOVA). **e-g.** Mitochondrial ROS levels (e), MitoTracker Green (ΔΨm-independent, f) and TMRM (ΔΨm-dependent, g) fluorescence intensity were quantified by flow cytometry in WT, KO, and KO+BNIP3L iN. Data are presented as mean ± SEM from one of three independent experiments. Statistical significance was assessed using one-way ANOVA followed by post hoc testing. ***P < 0.001, **P < 0.01, and ns, not significant. **h.** Representative confocal microscopy images of Pk-mito (red) and Hoechst (blue) double-stained WT, KO and KO+BNIP3L iN. Scale bar, 10 µm. **i-j.** Representative plot of oxygen consumption rate over time and timeline of injections with Oligomycin and Rotenone/Antimycin A (i) or FCCP and Rotenone/Antimycin A (j) as indicated. **k-m.** Basal oxygen consumption rate (OCR) (k), maximal OCR (l) and spare respiration capacity (m) calculated from the Seahorse assay shown above, in WT, KO, and KO+BNIP3L iN. Data are presented as mean ± SEM. Statistical significance was determined using one-way ANOVA. *P < 0.05; ns, not significant.

We next asked whether BNIP3L is required for neuronal mitophagy under basal conditions and in response to AD-associated stress. Mitophagy was quantified using the pH-sensitive mt-Keima reporter^30^ in WT, *BNIP3L* KO and BNIP3L-rescued neurons under basal conditions and following oligomeric Aβ42 exposure. *BNIP3L* KO significantly reduced basal mitophagy compared with WT neurons, supporting a role for BNIP3L in maintaining neuronal mitochondrial quality control (Fig. 2b). Oligomeric Aβ42 markedly increased mitophagy in WT neurons, whereas this response was largely abolished in *BNIP3*L KO neurons (Fig. 2b). Re-expression of BNIP3L restored both basal and Aβ42-induced mitophagy to levels comparable to WT neurons, demonstrating that the defect resulted specifically from BNIP3L loss (Fig. 2b). Together, these findings establish BNIP3L as a key mediator of basal neuronal mitophagy and the adaptive mitophagic response to Aβ-associated stress.

To determine whether this function is conserved *in vivo*, we examined DCT-1, the *C. elegans* orthologue of BNIP3L^23^. Using the pan-neuronal mtRosella mitophagy reporter, loss of *dct-1* significantly increased the GFP:DsRed fluorescence ratio, indicating reduced basal neuronal mitophagy compared with WT animals (Fig. 2c, d). Aβ_1-42_-expressing animals similarly exhibited reduced neuronal mitophagy, which was not further decreased by DCT-1 loss (Fig. 2c, d). These findings identify DCT-1 as a regulator of basal neuronal mitophagy *in vivo* and support an evolutionarily conserved role for the BNIP3L/DCT-1 pathway in neuronal mitochondrial turnover.

We next asked whether defective BNIP3L-dependent mitochondrial clearance compromises the quality of the neuronal mitochondrial pool. Ultrastructural analysis revealed an increased mitochondrial number in *BNIP3L* KO neurons (Fig. S2f), accompanied by reduced mitochondrial size (Fig. S2g) and shortened cristae (Fig. S2h), indicating the accumulation of smaller mitochondrial in *BNIP3L* KO neurons. Consistent with impaired mitochondrial turnover (Fig.2 a, b and Fig. S2c), BNIP3L-deficient neurons displayed increased mitochondrial ROS (Fig. 2e) and mitochondrial mass (Fig. 2f) together with reduced mitochondrial membrane potential (Fig. 2h). Re-expression of BNIP3L substantially restored these phenotypes linking defective mitochondrial quality to loss of BNIP3L (Fig. 2e–h). Thus, impaired BNIP3L-dependent mitophagy results in the accumulation of structurally and functionally compromised mitochondria in neurons.

Therefore, we next determined whether the accumulation of dysfunctional mitochondria limits neuronal bioenergetic capacity. Seahorse extracellular flux analysis revealed that basal respiration was largely preserved in BNIP3L KO neurons, whereas maximal respiration and spare respiratory capacity were reduced (Fig. 2i–m). BNIP3L re-expression rescued maximal respiration and respiratory reserve, indicating that BNIP3L becomes particularly important when neurons are challenged to increase mitochondrial energy production.

Together, these findings identify BNIP3L/DCT-1-dependent mitophagy as a conserved neuronal mitochondrial quality control pathway that supports both constitutive mitophagy and the maintenance of a functionally competent mitochondrial pool. Loss of this pathway permits the accumulation of damaged mitochondria and compromises maximal mitochondrial respiratory capacity required to meet increased energetic demand.

### DCT-1-dependent mitophagy preserves neuronal mitochondrial homeostasis *in vivo*

Having established that BNIP3L is required for neuronal mitophagy and mitochondrial homeostasis in human neurons, we next examined the consequences of disrupting this pathway *in vivo*. We used wild type and *dct-1(luc194)* mutant nematodes pan-neuronally expressing the mitochondria-targeted mKate2 fluorescent reporter (p*_rgef-1_*TOMM-20::mKate2) to visualize and quantify neuronal mitochondria. Loss of *dct-1* significantly increased mitochondrial mass in the head region of both wild-type and Aβ_1-42_-expressing animals at day 5 of adulthood (Fig. 3a, b), with similar effects observed at day 8 (Fig. S3a, b). Increased mitochondrial mass was also observed following *dct-1* loss in animals expressing pan-neuronal wild-type human Tau (EVL1654) (Fig. 3c, d and Fig. S3c, d). Thus, DCT-1 restricts neuronal mitochondrial accumulation under basal conditions and in the presence of AD-associated proteotoxic stress.

**Figure 3.**
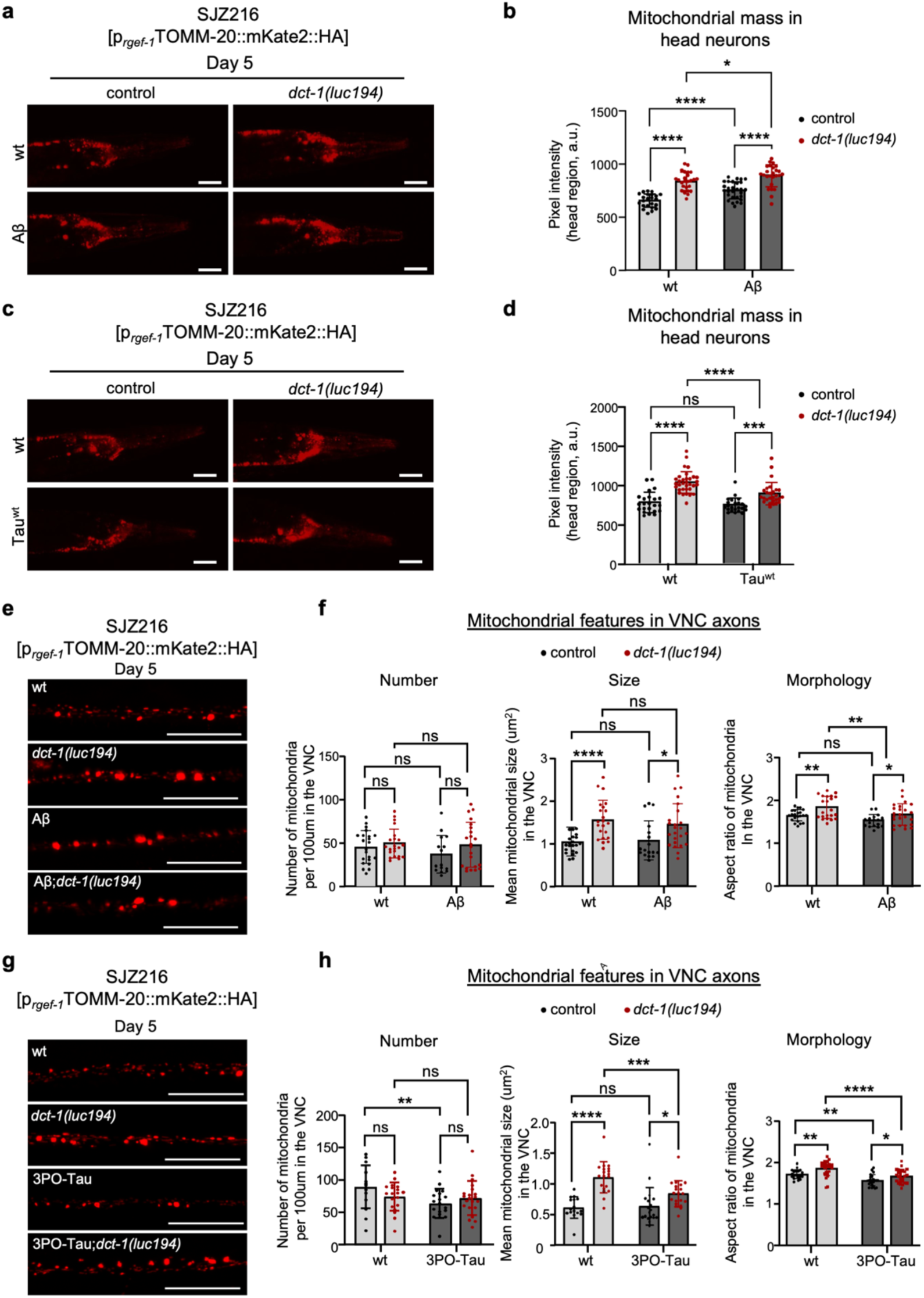
Loss of dct-1 (The homologue of BNIP3/BNIP3L in nematodes) alters mitochondrial content and axonal mitochondrial organization in *C. elegans*. **a.** Representative fluorescence images of the head region of 5-day-old adult *C. elegans* expressing the pan-neuronal mitochondrial reporter p*_rgef-1_*TOMM-20::mKate2::HA (SJZ216 strain in wild-type and Aβ-expressing (GRU102) backgrounds under control or *dct-1(luc194)* mutant conditions. Scale bar, 50 μm. **b.** Quantification of neuronal mitochondrial content based on the fluorescence intensity of the pan-neuronal mitochondrial reporter (p*_rgef-1_*:TOMM-20::mKate2::HA) in the head region of 5-day-old adult *C. elegans* (data presented as mean ± SD; * p < 0.05, ****p < 0.0001; one-way ANOVA). **c.** Representative fluorescence images of the head region of 5-day-old adult nematodes, expressing the pan-neuronal mitochondrial reporter p*_rgef-1_*TOMM-20::mKate2::HA (SJZ216 strain) in wild-type and Tau^wt^-expressing (EVL1654) backgrounds under control or *dct-1(luc194)* mutant conditions. Scale bar, 50 μm. **d.** Quantification of neuronal mitochondrial content based on the fluorescence intensity of the pan-neuronal mitochondrial reporter (p*_rgef-1_*:TOMM-20::mKate2::HA) in the head region of 5-day-old adult *C. elegans* (data presented as mean ± SD; * p < 0.05, ****p < 0.0001; one-way ANOVA). **e.** Representative fluorescence images of axonal mitochondria in 5-day-old adult *C. elegans* expressing the pan-neuronal mitochondrial reporter p*_rgef-1_*TOMM-20::mKate2::HA (SJZ216 strain) in wild-type and Aβ-expressing (GRU102) backgrounds under control or *dct-1(luc194)* mutant conditions. Scale bar, 20 μm. **f.** Quantification of axonal mitochondrial abundance (number of mitochondria per 100um axon length), mitochondrial size and morphology (aspect ratio) in the indicated genotypes (data presented as mean ± SD; ns p > 0.05, * p < 0.05, ****p < 0.0001; one-way ANOVA). **g.** Representative fluorescence images of axonal mitochondria in 5-day-old adult *C. elegans* expressing the pan-neuronal mitochondrial reporter p*_rgef-1_*TOMM-20::mKate2::HA (SJZ216 strain) in wild-type and 3POTau-expressing (EVL1372) backgrounds under control or *dct-1(luc194)* mutant conditions. Scale bar, 20 μm. **h.** Quantification of axonal mitochondrial abundance (number of mitochondria per 100um axon length), mitochondrial size and morphology (aspect ratio) in the indicated genotypes (data presented as mean ± SD; ns p > 0.05, * p < 0.05, ** p < 0.01, *** p < 0.001, ****p < 0.0001; one-way ANOVA).

We next asked whether the increased neuronal mass was accompanied by changes in mitochondrial abundance or morphology within individual axons. Therefore, we quantified mitochondrial number, size and shape in ventral nerve cord (VNC) axons. Despite the elevation in overall neuronal mitochondrial mass, *dct-1* loss had little effect on axonal mitochondrial number in wild-type or Aβ_1-42_-expressing nematodes. Instead, DCT-1 deficiency increased mitochondrial size at both day 5 and day 8 of adulthood (Fig. 3e, f and Fig. S3e, f). Mitochondrial aspect ratio, a measure of mitochondrial elongation, was also increased in both wild-type and Aβ_1-42_-expressing animals across ages. Thus, loss of DCT-1 leads predominantly to the accumulation of enlarged and elongated mitochondria within neuronal processes rather than simply increasing mitochondrial number.

We next investigated whether impaired mitochondrial quality control caused by DCT-1 deficiency modifies neuronal vulnerability to Tau proteotoxicity. We analyzed animals pan-neuronally expressing aggregation-prone 3PO-Tau (EVL1372) and quantified mitochondrial number, size and shape in VNC axons. At day 5, 3PO-Tau animals contained fewer axonal mitochondria than wild-type animals, with no further reduction following dct-1 loss (Fig. 3g, h). By day 8, however, the effect of DCT-1 deficiency differed according to the presence of Tau. Loss of DCT-1 reduced mitochondrial number in the wild-type background but increased mitochondrial number in 3PO-Tau nematodes (Fig. S3g, h). These findings indicate that Tau pathology modifies the mitochondrial morphological phenotype caused by dct-1 deficiency, with a relative shift from mitochondrial enlargement toward increased elongation that becomes more pronounced with age.

Tau also modified the effects of DCT-1 impairment on mitochondrial morphology. At day 5, *dct-1* loss markedly increased mitochondrial size in wild-type animals, whereas this effect was less pronounced in 3PO-Tau animals (Fig. 3g, h). This difference was also evident at day 8, when the mitochondrial enlargement associated with DCT-deficiency was attenuated in the 3PO-Tau background (Fig. S3g, h). By contrast, mitochondrial elongation, assessed by aspect ratio, was increased upon DCT-1 loss in 3PO-Tau nematodes at both ages and became particularly pronounced at day 8 (Fig. S3h). These observations show that Tau pathology modifies the remodeling caused by DCT-1 deficiency, with the interaction becoming more evident during aging.

Together, these findings demonstrate that DCT-1 is required to maintain neuronal mitochondrial homeostasis *in vivo*. Loss of DCT-1 leads to neuronal mitochondrial accumulation and alters mitochondrial size and shape within axons, consistent with defective mitochondrial turnover. Importantly, these consequences are influenced by both age and proteotoxic content, with Aβ and Tau differentially modifying the mitochondrial changes caused by DCT-1 deficiency.

### BNIP3L/DCT-1-dependent mitophagy restrains Tau and Aβ pathology across experimental systems

Having established that BNIP3L deficiency disrupts neuronal mitophagy and mitochondrial homeostasis, we next asked whether loss of this pathway promotes AD-associated protein pathology. In human iPSC-derived neurons, BNIP3L deficiency increased AD-associated Tau phosphorylation, whereas re-expression of BNIP3L reduced phospho-Tau towards control levels (Fig. 4a, b). By contrast, BNIP3L loss had minimal effects on Aβ42/Aβ40 and Aβ42/Aβ38 ratios under basal conditions (Fig. 4c). Following exposure to oligomeric Aβ42, however, BNIP3L KO neurons exhibited an increased Aβ42/Aβ40 ratio, which was partially restored by BNIP3L re-expression (Fig. 4d). Thus, BNIP3L deficiency promotes pathological Tau phosphorylation and alters Aβ homeostasis particularly under proteotoxic stress.

**Figure 4.**
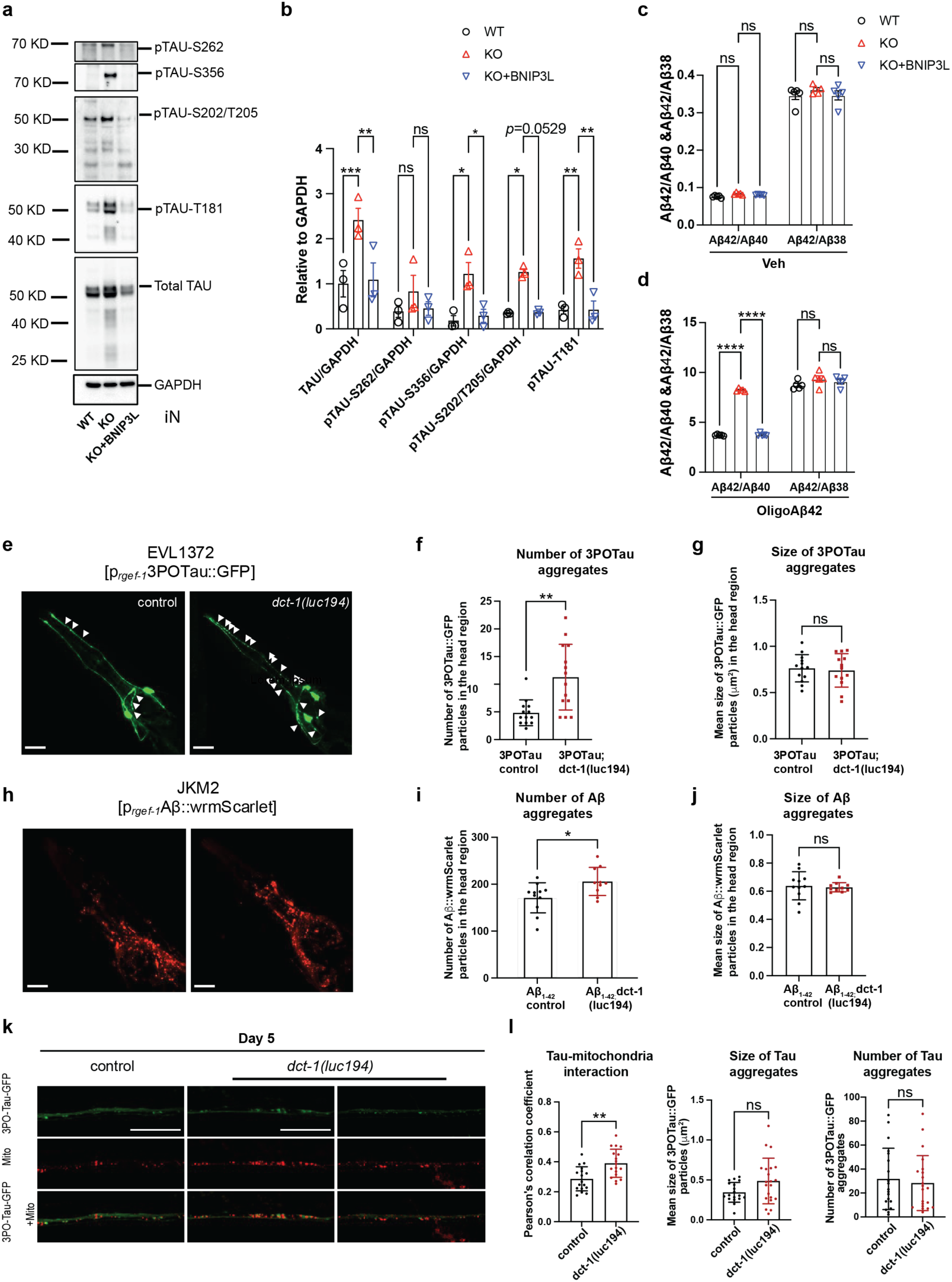
Loss of BNIP3L/dct-1 exacerbates tau and Aβ pathogenesis in nematodes and iPSC-derived neurons, while BNIP3L overexpression mitigates the pathogenesis in iPSC-derived neurons. **a.** Representative image of western blot data shows different AD-related TAU phosphorylation changes in WT, KO and KO+BNIP3L iN. **b.** Quantification of western blot data from three independent batches of experiments showing different AD-related TAU phosphorylation changes in WT, KO and KO+BNIP3L iN. **c-d:** Quantification of the Aβ42/Aβ40, Aβ42/Aβ38, and Aβ42/Aβ43 ratios in WT, BNIP3L knockout (KO), and KO+BNIP3L iN-cultured conditioned media under basal conditions (c) and following oligomeric Aβ42 treatment (Aβ42; 2.5 μM, 24 h) (d). The V-PLEX Plus Aβ Peptide Panel 1 (6E10) Kit (K15200G-1, MSD) was used to measure the Aβ42/Aβ40, Aβ42/Aβ38, and Aβ42/Aβ43 ratios. Data are presented as mean ± SEM. Statistical significance was determined using two-way ANOVA followed by Šidák’s multiple-comparisons test. ****P < 0.0001; ns, not significant. **e.** Representative images of the head region of 5-day adult nematodes expressing pan-neuronally (rgef-1 promoter) 3POTau::GFP transgene (EVL1372 strain) under control or dct-1(luc194) mutant conditions. Arrows indicate 3POTau::GFP-positive particles. Scale bar, 20 μm. **f.** Quantification of the number of 3POTau::GFP positive particles in the head region of the indicated genotypes (data presented as mean ± SD; * p < 0.05; Unpaired t-test). **g.** Quantification of the size of 3POTau::GFP positive particles in the head region of the indicated genotypes (data presented as mean ± SD; ns p > 0.05; Unpaired t-test). **h.** Representative images of the head region of 5-day adult nematodes expressing pan-neuronally (rgef-1 promoter) Αβ*_1-42_*::wrmScarlet transgene under control or *dct-1(luc194)* mutant conditions. Scale bar, 20 μm. **i.** Quantification of the number of Αβ*_1-42_*::wrmScarlet positive particles in the head region of the indicated genotypes (data presented as mean ± SD; * p < 0.05; Unpaired t-test). **j.** Quantification of the size of Αβ*_1-42_*::wrmScarlet positive particles in the head region of the indicated genotypes (data presented as mean ± SD; ns p > 0.05; Unpaired t-test). **k.** Representative images of the axonal region of 5-day adult nematodes co-expressing the 3POTau::GFP and the TOMM-20::mKate2::HA transgenes (EVL1372;SJZ216 strain) under control or *dct-1(luc194)* mutant conditions. **l.** Quantification of the Pearson’s correlation coefficient (PCC), the number of 3POTau::GFP positive particles and the size of 3POTau::GFP positive particles between 3POTau::GFP and the mitochondrial marker TOMM-20::mKate2::HA in the head region of 5-day-old adult nematodes under control or *dct-1(luc194)* mutant conditions. PCC was calculated to assess the degree of Tau–mitochondria colocalization. (data presented as mean ± SD; ** p < 0.01; Unpaired t-test).

We next asked whether loss of the conserved BNIP3L/DCT-1 pathway similarly affects protein aggregation-prone 3PO-Tau or Aβ_1-42_ pan-neuronally and quantified protein aggregates in the head region of *C. elegans*. Loss of DCT-1 significantly elevated the number of 3PO-Tau-positive aggregates in young adult nematodes without altering aggregate size (Fig. 4e–g). Similarly, DCT-1 deficiency increased neuronal Aβ aggregation *in vivo* (Fig. 4h–j). Thus, loss of DCT-1 promotes the accumulation of both Tau and Aβ aggregates in neurons *in vivo*, extending the relationship between BNIP3L/DCT-1-dependent mitochondrial quality control and AD-associated proteinopathy across species.

Because Tau can associate with mitochondria and perturb mitochondrial function^31^, we next asked whether DCT-1 deficiency influences the association of Tau with neuronal mitochondria *in vivo*. Loss of DCT-1 increased Tau-mitochondria association at both day 5 and day 8 of adulthood (Fig. 4k, l and Fig. S4a, b). This increase was already detectable at day 5 and became more pronounced with age, indicating that impaired DCT-1-dependent mitochondrial quality control is accompanied by increased association of Tau with the neuronal mitochondrial network. Whether this increased association contributes to Tau aggregation or instead reflects preferential accumulation of Tau at dysfunctional mitochondria remains to be determined.

Having found that loss of BNIP3L/DCT-1 exacerbates Tau pathology, we next examined whether increasing BNIP3Lexpression is sufficient to suppress pathological Tau accumulation. Primary mouse cortical neurons were exposed to Tau seeds to induce pathological Tau aggregation and transduced with lentivirus expressing human BNIP3L (LV-BNIP3L). BNIP3L overexpression markedly reduced seeded Tau aggregation (Fig. 5a,b and Fig. S5a). Immunofluorescence analysis confirmed robust BNIP3L expression and further revealed reduced AT8-positive phosphorylated Tau (pSer202/Thr205) aggregates upon BNIP3L overexpression (Fig. 5c,d). Thus, increasing BNIP3L is sufficient to suppress seeded Tau pathology in primary neurons.

**Figure 5.**
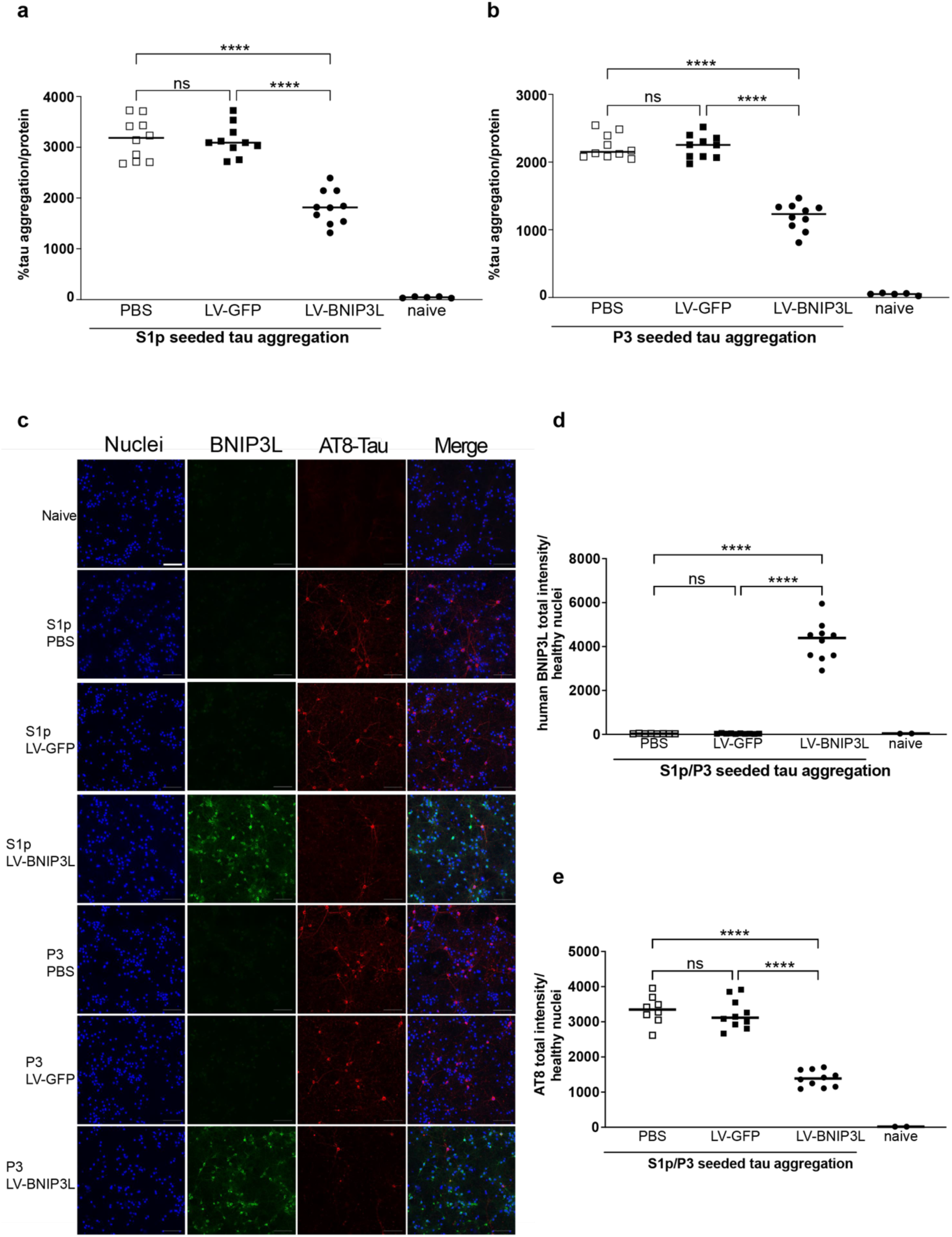
BNIP3L overexpression mitigates tau pathology in mouse cortical neurons. a-b. S1p seeded tau aggregation (a) and P3 seeded tau aggregation (b) in primary mouse cortical neurons from rTg4510 mice under different conditions, including PBS, LV-GFP, LV-BNIP3L, and naïve controls. LV-GFP, lentiviral GFP; LV-BNIP3L, lentiviral BNIP3L. Naïve controls were not exposed to tau seeds. Data are presented as the percentage seeded Tau aggregation normalized to total protein as mean ± SD. Statistical significance was determined using Dunnett’s T3 multiple-comparisons test. \*\*\*\**P* < 0.0001; ns, not significant. **c.** Representative confocal images (20× objective) of nuclei stained with Hoechst (in blue), of BNIP3L immunoreactivity (in green), and of seeded Tau aggregation detected by AT8-Tau (pTau-S202/T205) immunoreactivity (in red) from primary mouse cortical neurons of rTg4510 mice seeded with S1p and P3 under different conditions, including PBS, LV-GFP, LV-BNIP3L, and naïve controls. Naïve controls were not exposed to tau seeds. Data are presented as the median. The scale bar corresponds to 100 μm. **d-e.** Quantification of human BNIP3L immunoreactivity (d) and AT8-Tau immunoreactivity (e) in primary mouse cortical neurons under different conditions, including PBS, LV-GFP, LV-BNIP3L, and naïve controls. Data are presented as total BNIP3L and AT8 intensity normalized to healthy nuclei as mean ± SD. Statistical significance was determined using Dunnett’s T3 multiple-comparisons test. ****P < 0.0001; ns, not significant.

Taken together, these findings provide bidirectional evidence linking the BNIP3L/DCT-1 pathway to AD-associated proteinopathy across human neurons, *C. elegans* and primary mouse neurons. Loss of BNIP3:/DCT-1 enhances Tau phosphorylation and aggregation and promotes Aβ pathology, whereas restoring or increasing BNIP3L suppresses pathological Tau accumulation. These findings identify BNIP3L/DCT-1 dependent mitophagy as a conserved pathway that restrains AD-associated proteotoxicity.

### BNIP3L is required for neuronal maturation and network function

Having established that BNIP3L deficiency disrupts mitochondrial homeostasis and promotes AD-associated pathology, we next examined how loss of BNIP3L affects neuronal function. We first performed quantitative proteomic profiling of WT, BNIP3L KO and BNIP3L-rescued human iPSCs-derived neurons to identify cellular pathways altered by BNIP3L deficiency and restored following its re-expression.

BNIP3L KO neurons showed broad proteomic remodeling involving proteins associated with mitochondria, lysosomes, amyloid-related processes and the cytoskeleton (Fig. 6a). Many of these alterations were partially reversed upon BNIP3L re-expression (Fig. 6a). Gene ontology analysis further revealed enrichment of mitochondrial and lysosomal compartments, synaptosomes, amyloid-related structures and intermediate filaments among proteins altered by BNIP3L deficiency (Fig. 6b). Notably, proteins reduced following BNIP3L restoration were enriched for mitochondrial components (Fig. 6c), whereas proteins increased following restoration were associated with glutamatergic synaptic signaling, vesicular trafficking and endosomal organization (Fig. S6b). These proteomic changes indicate that BNIP3L deficiency extends beyond mitochondrial homeostasis and is associated with alterations in pathways important for neuronal structure and synaptic function.

**Figure 6.**
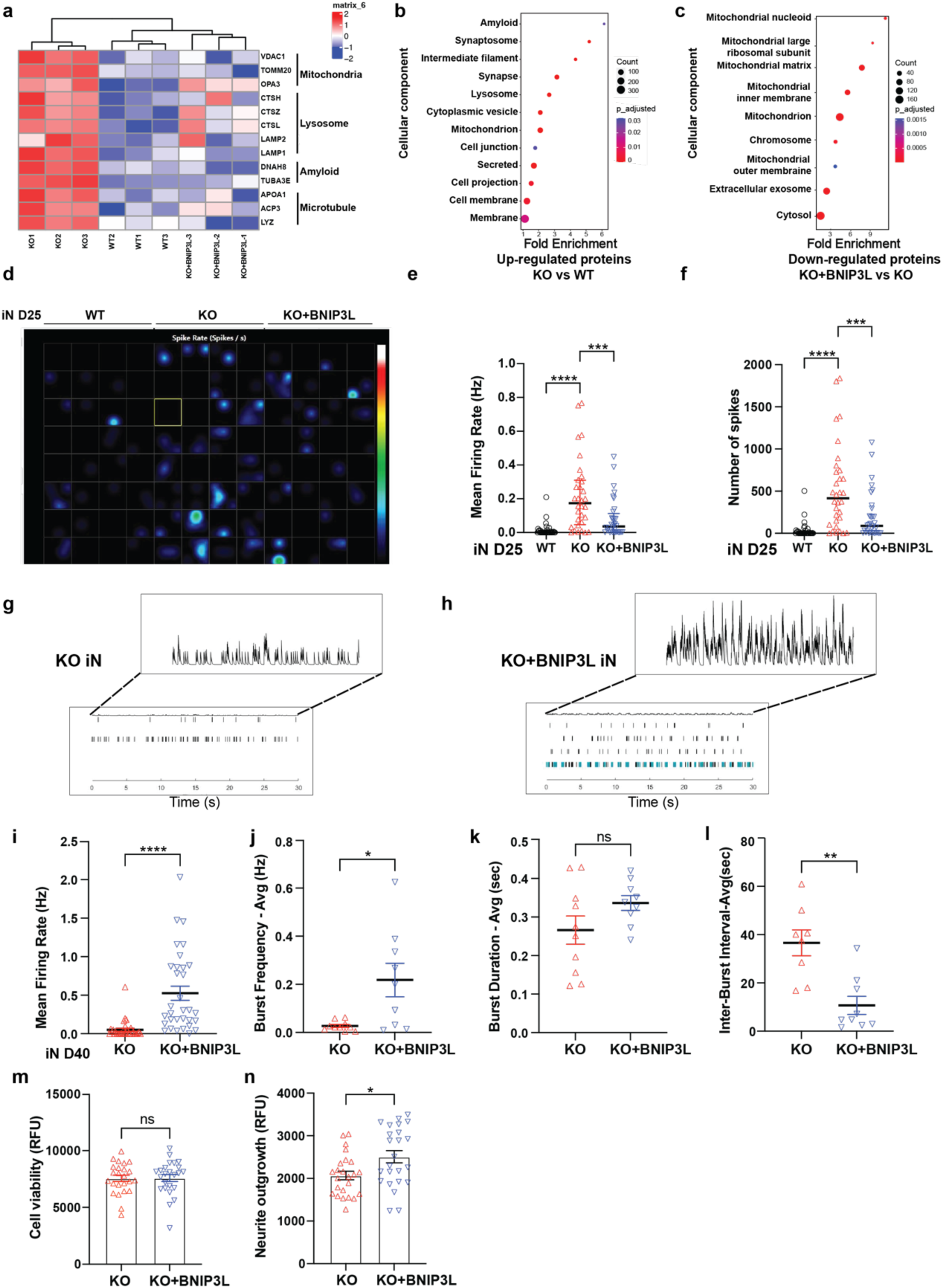
Proteomic and Electrophysiological Analyses Reveal a Neuroprotective Role for BNIP3L in iPSC-Derived Neurons. **a.** Heatmap showing dysregulated protein expression profiles in iN, including mitochondrial-related proteins (VDAC1, TOMM20, OPA3), lysosome-related proteins (LAMP1, LAMP2, CTSH, CTSZ, and CTSL), amyloid-related proteins (TUBA3E), microtubule-related proteins (APOA1, ACP3, and LYZ). Red and blue indicate increased and decreased protein expression levels, respectively, based on Z-scores. **b. b-c.** The gene ontology (GO) cellular component analysis shows the changes associated with up-regulated proteins (b) between KO iN and WT iN and down-regulated proteins (c) between KO iN overexpressed with BNIP3L iN and KO iN. The GO biological processes were organized by Fold enrichment and p_adjusted. **c.** d. The heatmap image showing the mean firing rate in in WT, KO and KO+BNIP3L iPSCs-derived neuron at differentiation day 25. **d.** e-f. Graph showing comparison of mean firing rate (e) and number of spikes (f) in WT, KO and KO+BNIP3L iPSCs-derived neuron at differentiation day 25. **e.** g-h. Representative raster plots of 30 seconds (sec) showing network activity of KO (g) and KO+BNIP3L iPSCs-differentiated neurons at differentiation day 40. **f.** i-l. Graph showing comparison of four MEAs, including mean firing rate (i), burst frequency (j), burst duration (k) and inter-burst interval (l) between KO and KO+BNIP3L iN at day40. Data are presented as mean ± SEM. Statistical analyses were conducted with two-tailed unpaired t-test. **** p < 0.0001, * p < 0.05, ns, no significant difference. **g.** m-n. Quantification of cell viability (m) and neurite outgrowth (n) in KO and KO+BNIP3L iN at day40. Data are presented as mean ± SEM. Statistical analyses were conducted with two-tailed unpaired t-test. * p < 0.05, ns, no significant difference.

Therefore, we asked whether these molecular changes are accompanied by altered neuronal network function. To follow the development of spontaneous neuronal activity over time, we performed longitudinal microelectrode array (MEA) recordings^32^. BNIP3L KO neurons initially exhibited a transient increase in firing activity during early stages of network development (days 25–27; Fig. 6d–f). This early increase was transient and was not maintained as the neuronal cultures matured (Fig. S6c–e). At later stages, BNIP3L KO cultures instead showed reduced mean firing rate, burst frequency and burst duration, together with prolonged intervals between burst, compared with BNIP3L-rescued neurons (Fig. 6g–l). Thus, BNIP3L deficiency alters the developmental trajectory of neuronal network activity, characterized by transient early hyperactivity followed by impaired network activity at later stages.

We next examined whether impaired network activity reflected neuronal loss or defective neuronal maturation. Neuronal viability was unchanged by BNIP3L deficiency (Fig. 6m), arguing against cell loss as the primary explanation for the reduced network activity. By contrast, neurite outgrowth was significantly reduced in BNIP3L KP neurons compared with BNIP3L-rescued neurons (Fig. 6n). Thus, BNIP3L deficient neurons remain viable but exhibit impaired structural development together with an inability to sustain normal network activity as they mature.

Together, these findings identify BNIP3L as an important determinant of neuronal maturation and network function. Loss of BNIP3L induces broad molecular remodeling, including changes in synaptic and neuronal pathways, and disrupts the normal development of neuronal activity without causing excessive neuronal loss. These findings connect defective mitochondrial quality control with impaired structural and functional neuronal maturation.

### DCT-1 supports learning and memory and cholinergic neurotransmission *in vivo*

Having demonstrated that BNIP3L deficiency compromises neuronal network function in human neurons, we next asked whether loss of the conserved BNIP3L orthologue DCT-1 results in functional neuronal deficits at the organismal level. Firstly, we assessed associative learning and memory in wild-type and AD nematode models in the presence or absence of *dct-1*. Loss of DCT-1 significantly impaired learning and memory in otherwise wild-type animals, as indicated by reduced avoidance of isoamyl alcohol following conditioning (Fig. 7a). Aβ_1-42_-expressing nematodes were already impaired compared to wild-type controls, and DCT-1 deficiency did not further worsen their behavioral performance (Fig. 7b). By contrast, 3PO-Tau expression alone did not significantly impair learning and memory, whereas DCT-1 loss in the 3PO-Tau background markedly reduced their performance (Fig. 7a). Thus, DCT-1 is required for normal associative learning and memory and become particularly important for preserving behavioral function in the presence of pathological Tau.

**Figure 7.**
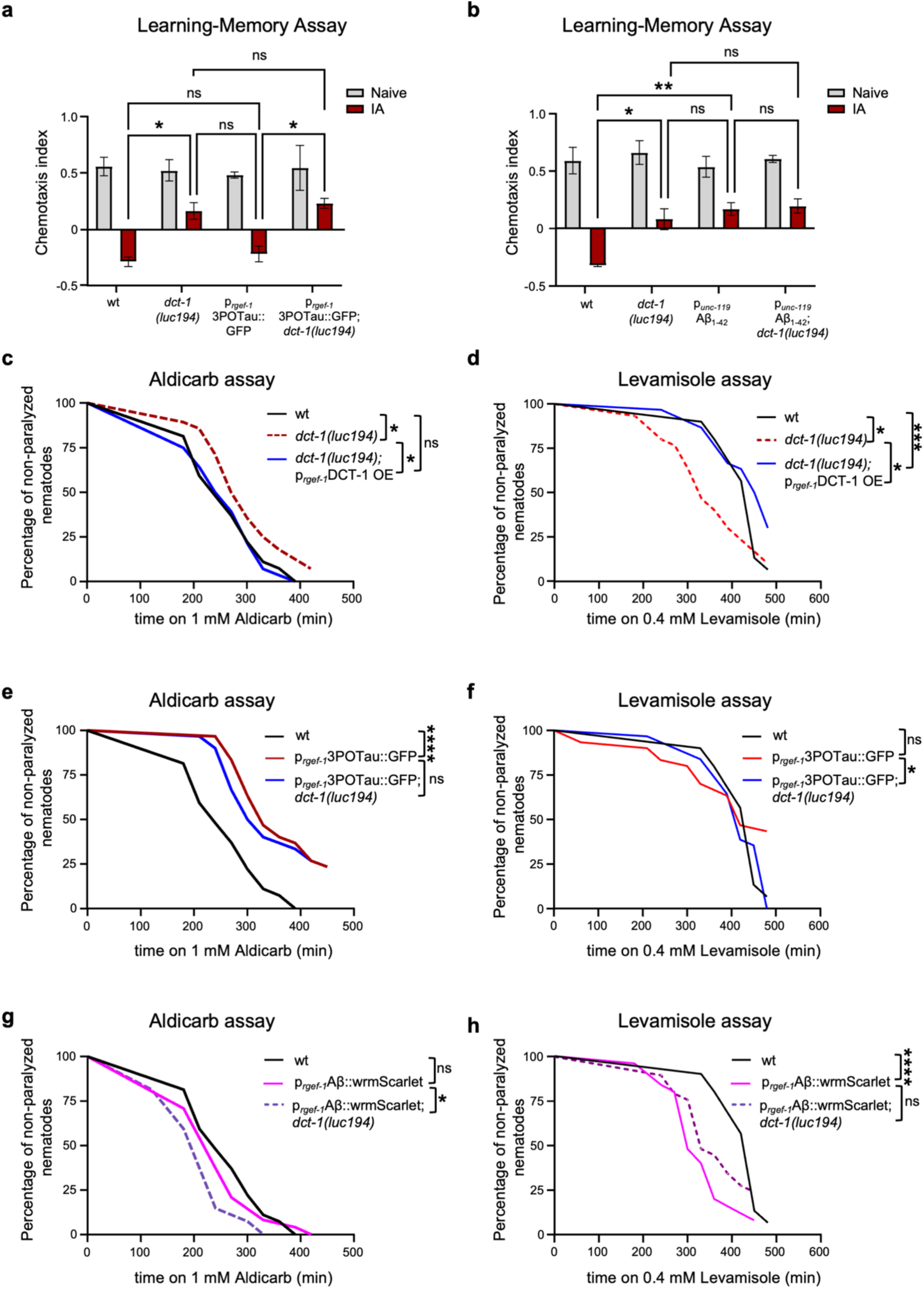
Loss of dct-1 decrease cognition and affect synaptic transmission in nematodes. **a.** Chemotaxis index of 1-day adult wt (N2) and 3POTau-expressing (EVL1372 strain) nematodes under control and *dct-1(luc194)* mutant conditions following associative learning with the chemo-attractant Isoamyl Alcohol (IA). The nematodes were conditioned for 90min on plates without bacteria either in the absence (Naive group) or presence (IA group) of IA and subsequently transferred to chemotaxis assay plates to evaluate their attraction to IA, as a measure of associative learning and memory, the nematodes were transferred to assay plates to assess their chemotaxis index to IA in order to check their learning and memory capacity (data presented as mean ± SEM from 3 biological repeats, ns p>0.05, p<0,05, one-way ANOVA). **b.** Chemotaxis index of 1-day adult wt (N2) and Αβ-expressing (GRU102 strain) nematodes under control and *dct-1(luc194)* mutant conditions following associative learning with the chemo-attractant Isoamyl Alcohol (IA). The nematodes were conditioned for 90min on plates without bacteria either in the absence (Naive group) or presence (IA group) of IA and subsequently transferred to chemotaxis assay plates to evaluate their attraction to IA, as a measure of associative learning and memory, the nematodes were transferred to assay plates to assess their chemotaxis index to IA in order to check their learning and memory capacity (data presented as mean ± SEM from 3 biological repeats, ns p>0.05, p<0,05, one-way ANOVA). **c. c–d.** Paralysis curves of 1-day-old adult wild-type (N2), dct-1(luc194) mutant, and dct-1(luc194);Prgef-1::DCT-1 OE rescue nematodes following exposure to 1 mM aldicarb (c) or 0.4 mM levamisole (d). **d.** e–f. Paralysis curves of 1-day-old adult wild-type (N2), 3PO-Tau-expressing (EVL1372), and 3PO-Tau-expressing;dct-1(luc194) mutant nematodes following exposure to 1 mM aldicarb (e) or 0.4 mM levamisole (f). **e.** g-h. Paralysis curves of 1-day-old adult wild-type (N2), Aβ-expressing (JKM2), and Aβ-expressing;dct-1(luc194) mutant nematodes following exposure to 1 mM aldicarb (g) or 0.4 mM levamisole (h). **f.** For c–h, differences between paralysis curves were analyzed using the log-rank (Mantel–Cox) test. ns, P > 0.05; *P < 0.05; **P < 0.01; ***P < 0.001; ****P < 0.0001.

We next examined whether these behavioral deficits are accompanied by altered synaptic transmission. We focused on cholinergic neurotransmission at the neuromuscular junction using complementary aldicarb and levamisole assays^33, 34^. Aldicarb inhibits acetylcholinesterase, leading to accumulation of acetylcholine at the synaptic cleft and progressive paralysis; resistance to aldicarb is therefore consistent with reduced presynaptic acetylcholine release. In contrast, levamisole directly activates postsynaptic nicotinic acetylcholine receptors and provides a complementary measure of postsynaptic responsiveness. Notably, *dct-1(luc194)* mutant nematodes displayed a modest shift toward aldicarb resistance, whereas pan-neuronal re-expression of DCT-1 restored aldicarb responses towards wild-type levels (Fig. 7c). These findings are consistent with reduced cholinergic output following loss of neuronal DCT-1 and, importantly show that restoring DCT-1 in neurons is sufficient to reverse this functional phenotype.

We next examined postsynaptic responsiveness using levamisole. DCT-1 deficient nematodes displayed increased sensitivity to levamisole-induced paralysis compared with wild-type animals, whereas neuronal DCT-1 re-expression shifted the response towards wild-type levels (Fig. 7d). Thus, loss of DCT-1 is associated with reduced presynaptic cholinergic output together with enhanced postsynaptic responsiveness., and both phenotypes are reversed towards normal by restoring DCT-1 expression in neurons. These opposing responses suggest that DCT-1 contributes to the coordinated regulation of cholinergic neuromuscular transmission, with enhanced postsynaptic responsiveness potentially representing an adaptation to reduced neuronal output.

Then, we examined whether Tau pathology alters the requirement for DCT-1 in cholinergic neurotransmission. 3PO-Tau-expressing animals displayed pronounced aldicarb resistance compared with wild-type controls (Fig. 7e), consistent with impaired presynaptic cholinergic transmission. Additionally, loss of DCT-1 did not further increase aldicarb resistance. In contrast, 3PO-Tau animals showed levamisole responses comparable to wild-type animals, whereas DCT-1 loss significantly altered the kinetics of levamisole-induced paralysis (Fig. 7f). Thus, Tau pathology is associated predominantly with impaired presynaptic cholinergic transmission, whereas DCT-1 deficiency additionally modifies postsynaptic responsiveness in the presence of pathological Tau.

A distinct pattern emerged in animals expressing neuronal Aβ_1-42_. Aβ_1–42_-expressing nematodes showed modest aldicarb hypersensitivity, which was further enhanced by DCT-1 deficiency (Fig. 7g). Aβ_1-42_ expression also caused pronounced levamisole hypersensitivity compared with wild-type controls, whereas DCT-1 deficiency partially attenuated this response (Fig. 7h). Thus, rather than uniformly worsening synaptic dysfunction, DCT-1 deficiency differentially modifies pre- and postsynaptic responses in the presence of Aβ.

Taken together, these findings demonstrate that neuronal DCT-1 supports associative learning and memory and contributes to the coordinated regulation of cholinergic neurotransmission *in vivo*. Importantly, neuronal re-expression of DCT-1 reverses the synaptic defects caused by DCT-1 deficiency, providing functional evidence that restoration of the BNIP3L/DCT-1 pathway can recover neuronal function. Under AD-associated proteotoxic conditions, however, the consequences of DCT-1 deficiency are context-dependent, revealing distinct functional interactions with Tau and Aβ pathology.

## Discussion

A central question in AD is how defective mitochondrial quality control contributes to proteinopathy and neuronal dysfunction. Here, we identify the BNIP3L/DCT-1-pathway as a conserved regulator of neuronal mitophagy that links mitochondrial homeostasis to AD-associated pathology and neuronal function. Loss of BNIP3L/DCT-1 impaired basal and stress-induced mitophagy, compromised mitochondrial integrity and respiratory capacity, enhanced Tau and Αβ pathology, and disrupted neuronal network, synaptic and behavioral function across human neurons and *C. elegans*. Conversely, BNIP3L restoration reversed mitochondrial and functional phenotypes, while increasing BNIP3L expression markedly suppressed seeded Tau aggregation in primary mouse neurons.

These findings extend current models of mitophagy dysfunction in AD by identifying receptor-mediated mitochondrial clearance as a critical determinant of neuronal mitochondrial quality, proteostasis and function (Fig. 8).

**Figure 8.**
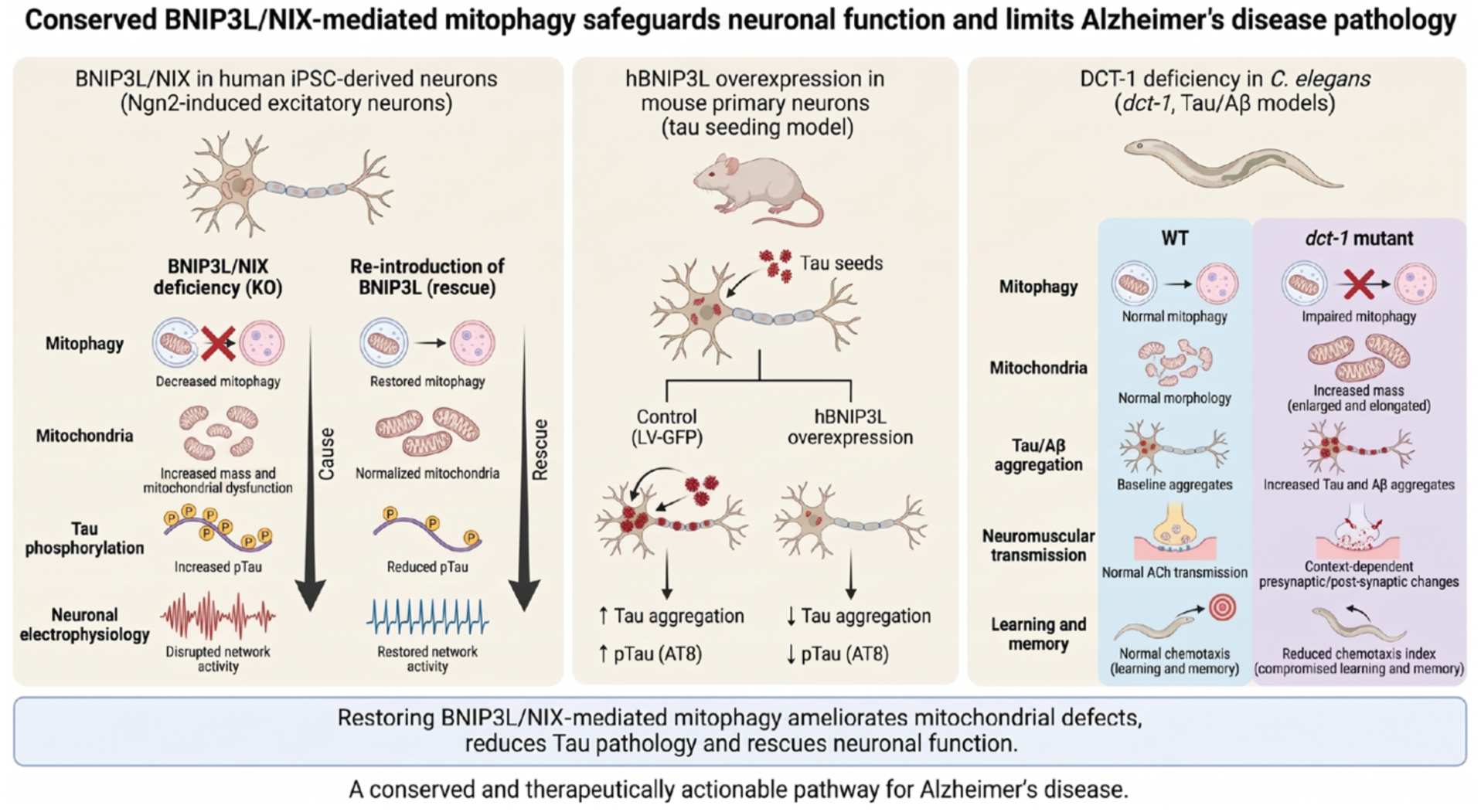
Model Figure. BNIP3L/NIX maintains neuronal mitochondrial homeostasis and function through mitophagy-mediated mitochondrial quality control. Under physiological conditions, BNIP3L promotes the clearance of damaged mitochondria, preserving mitochondrial integrity, neuronal activity, and cognitive function while limiting tau and Aβ pathology. Loss of BNIP3L impairs mitophagy, leading to mitochondrial defects, ROS accumulation, tau hyperphosphorylation, enhanced Aβ pathology, and neuronal dysfunction. Conversely, restoration of BNIP3L improves mitochondrial function, while overexpression of human BNIP3L suppresses seeded tau aggregation in primary mouse neurons, demonstrating that enhancement of BNIP3L activity can mitigate pathogenic tau propagation. In *C. elegans*, loss of DCT-1 exacerbates tau- and Aβ-induced deficits in learning, memory, and locomotion, supporting a critical role for BNIP3L-mediated mitophagy in maintaining neuronal and cognitive health. Together, findings from human iPSC-derived neurons, primary mouse neurons and nematode models establish BNIP3L/NIX-mediated mitophagy as a conserved pathway linking mitochondrial homeostasis to tau and Aβ pathology and support BNIP3L as a potential therapeutic target for Alzheimer’s disease and related tauopathies.

BNIP3L-mediated mitophagy may have a broader role in neuronal homeostasis than previously recognized. In *C. elegans*, DCT-1 functions as a mitophagy coordinator that couples mitochondrial clearance to broader programs of mitochondrial biogenesis^14^, while neuronal BNIP3 induction promotes mitochondrial homeostasis and longevity in *D. melanogaster*^20^. BNIP3L-dependent mitophagy also supports retinal ganglion cell differentiation by facilitating mitochondrial degradation and metabolic remodeling^22^. In our study, BNIP3L expression increased markedly during neuronal differentiation, consistent with an increased requirement for mitochondrial quality control as neurons acquire their mature metabolic and functional properties. Moreover, BNIP3L abundance in the human brain was inversely associated with neuropathological and clinical measures of AD severity. Together, these observations suggest that BNIP3L-dependent mitochondrial quality control is engaged during neuronal maturation and may become increasingly important for preserving neuronal homeostasis and survival in the context of AD pathogenesis.

Our findings also broaden the mechanistic framework of neuronal mitophagy beyond the extensively studied PINK1-PARKIN pathway. Although PINK1-PARKIN mediated mitophagy is prominently engaged following mitochondrial damage, basal mitophagy can persist independently of PINK1 or PARKIN in several tissues and neuronal populations *in vivo*^17^. We found that BNIP3L loss reduced basal mitophagy in human neurons, while DCT-1 deficiency impaired neuronal mitophagy *in vivo,* identifying receptor-mediated mitochondrial clearance as an important component of constitutive neuronal mitochondrial turnover. Importantly, Aβ42 increased mitophagy in control neurons, whereas this adaptive response was largely abolished in BNIP3L-deficient neurons. Thus, BNIP3L-depdendent mitophagy contributes not only to basal mitochondrial maintenance but also to the ability of neurons to increase mitochondrial clearance in response to AD-associated proteotoxic stress.

The mitochondrial consequences of BNIP3L/DCT-1 deficiency further support a pivotal role for this pathway in maintaining a functional neuronal mitochondrial pool. BNIP3L deficient human neurons accumulated mitochondria with reduced membrane potential, ultrastructural abnormalities and elevated ROS. Despite relatively preserved basal respiration, maximal respiration and spare respiratory capacity were impaired. This distinction suggests that defective mitochondrial clearance does not cause immediate bioenergetic collapse but instead reduces the capacity of neurons to handle their enhanced energetic demands. DCT-1 deficiency similarly increased neuronal mitochondrial content and altered mitochondrial morphology *in vivo*, supporting a conserved requirement for the BNIP3L/DCT-1 pathway in neuronal homeostasis. These mitochondrial phenotypes were further influenced by age and proteotoxic content, particularly in Tau-expressing nematodes. A very recent study uncovered that loss of Tau/PTL-1 promotes a mitofusin-dependent pro-fusion state and alters mitochondrial bioenergetics and stress adaptation^31^. Together with the increased Tau-mitochondria association observed upon DCT-1 deficiency, these findings point to a reciprocal relationship between mitochondrial quality control and Tau-dependent mitochondrial remodeling. Whether mitochondrial elongation associated with DCT-1 deficiency results from altered fusion-fission dynamics, persistence of mitochondria normally targeted for removal, or a combination of these processes, remains to be determined.

A major conceptual advance in this study is the identification of BNIP3L/DCT-1 pathway as a regulator of pathological Tau accumulation. DCT-1 loss increased neuronal Tau aggregation *in vivo* in *C. elegans*, whereas BNIP3L deficiency enhanced disease-associated Tau phosphorylation in human neurons. Conversely, BNIP3L restoration reduced Tau phosphorylation and markedly suppressed seeded Tau aggregation in primary mouse neurons. DCT-1 loss also increased the association of Tau with neuronal mitochondria, an effect that became more pronounced with age. Whether this increased association contributes to subsequent Tau aggregation, reflects preferential recruitment of Tau to dysfunctional mitochondria, or results from the persistence of mitochondria that would normally be removed remains unresolved. Nevertheless, the reciprocal effects of BNIP3L/DCT-1 loss and restoration across experimental systems provide bidirectional evidence linking this mitochondrial quality control pathway to pathological Tau accumulation.

BNIP3L/DCT-1 also influenced Aβ pathology, although these effects were more context dependent. DCT-1 deficiency increased Aβ aggregation *in vivo*, whereas BNIP3L deficiency had little effect on Aβ42/Aβ40 or Aβ42/Aβ38 ratios in human neurons under basal conditions. Following Aβ42 challenge, however, BNIP3L deficient neurons showed an increased Aβ42/Aβ40 ratio that was partially restored by BNIP3L re-expression. Together with the attenuated BNIP3L response to Aβ42 observed in sAD neurons, these findings suggest that BNIP3L dependent mitochondrial quality control may become particularly important when neurons are challenged by Αβ-associated proteotoxic stress. The distinct functional responses observed following DCT-1 loss in Tau- and Aβ-expressing nematodes further indicate that disruption of this pathway does not uniformly amplify proteotoxicity but instead interacts with the underlying pathological context.

Importantly, the consequences of BNIP3L/DCT-1 deficiency extended beyond mitochondrial dysfunction and proteinopathy to neuronal function. BNIP3L-deficient human neurons showed an altered developmental trajectory of network activity, characterized by transient early hyperactivity followed by reduced firing and bursting as cultures matured. These abnormalities occurred without detectable loss of neuronal viability but were accompanied by reduced neurite outgrowth, suggesting impaired neuronal maturation rather than non-specific cell loss. In *C. elegans*, DCT-1 deficiency impaired associative learning and memory and altered neurotransmission. Moreover, pan-neuro al restoration of DCT-1 shifted both aldicarb and levamisole responses towards wild-type levels, supporting a neuronal requirement for DCT-1 in maintaining coordinated synaptic function. Notably, the functional consequences of DCT-1 deficiency differed between Tau- and Aβ-expressing nematodes, indicating that mitophagy dysfunction interacts with the underlying proteotoxic environment rather than uniformly exacerbating neuronal dysfunction. Together, these findings connect mitochondrial quality control to neuronal function across structural, network, synaptic and behavioral levels.

Our findings have established how defective mitochondrial clearance is mechanistically coupled to Tau pathology and neuronal dysfunction. The accumulation of ROS-producing mitochondria could alter Tau phosphorylation through stress-responsive kinases such as GSK3β, p38 MAPK, JNK and CDK5^35, 36^, while altered endolysosomal trafficking^4, 37^ or other mitochondrial stress responses may influence Tau localization, aggregation or clearance when BNIP3L-dependent mitophagy is impaired. Increased Tau association with mitochondria provides another potential point of interaction between defective mitochondrial quality control and Tau pathology. An important next step will be to determine whether these downstream effects specifically require the mitophagy function of BNIP3L or involve additionally functions of the protein. Such studies should help define the molecular mechanisms connecting receptor mediated mitophagy to neuronal proteostasis.

Importantly, we identify BNIPR3L/DCT-1-dependent mitophagy as a conserved neuronal mitochondrial quality control pathway linking mitophagy, proteostasis and neuronal function. The ability of BNIP3/DCT-1 restoration to recover mitochondrial and neuronal function and suppress pathological Tau accumulation highlights receptor-mediated mitophagy as a potential therapeutic strategy for AD and related tauopathies.

## References

1. Abdulkhaliq, A.A. et al. Amyloid-beta and Tau in Alzheimer’s disease: pathogenesis, mechanisms, and interplay. Cell Death Dis 17, 21 (2026).

2. Zhang, J., Zhang, Y., Wang, J., Xia, Y., Zhang, J. & Chen, L. Recent advances in Alzheimer’s disease: Mechanisms, clinical trials and new drug development strategies. Signal Transduct Target Ther 9, 211 (2024).

3. Chan, D.C. Mitochondrial Dynamics and Its Involvement in Disease. Annu Rev Pathol 15, 235–259 (2020).

4. Du, H., Guo, L., Yan, S., Sosunov, A.A., McKhann, G.M. & Yan, S.S. Early deficits in synaptic mitochondria in an Alzheimer’s disease mouse model. Proc Natl Acad Sci U S A 107, 18670–18675 (2010).

5. Giorgi, C., Marchi, S. & Pinton, P. The machineries, regulation and cellular functions of mitochondrial calcium. Nat Rev Mol Cell Biol 19, 713–730 (2018).

6. Klemmensen, M.M., Borrowman, S.H., Pearce, C., Pyles, B. & Chandra, B. Mitochondrial dysfunction in neurodegenerative disorders. Neurotherapeutics 21, e00292 (2024).

7. Li, Y., Berliocchi, L., Li, Z. & Rasmussen, L.J. Interactions between mitochondrial dysfunction and other hallmarks of aging: Paving a path toward interventions that promote healthy old age. Aging Cell 23, e13942 (2024).

8. Swerdlow, R.H. Mitochondria and Mitochondrial Cascades in Alzheimer’s Disease. J Alzheimers Dis 62, 1403–1416 (2018).

9. Yao, J., Irwin, R.W., Zhao, L., Nilsen, J., Hamilton, R.T. & Brinton, R.D. Mitochondrial bioenergetic deficit precedes Alzheimer’s pathology in female mouse model of Alzheimer’s disease. Proc Natl Acad Sci U S A 106, 14670–14675 (2009).

10. Castillo-Casana, Y., Arias, C. & Coria, R. Tau-Mitochondria Interactions in Neurodegeneration: Mechanisms and Therapeutic Potential. Cell Mol Neurobiol 46, 3 (2025).

11. Onishi, M., Yamano, K., Sato, M., Matsuda, N. & Okamoto, K. Molecular mechanisms and physiological functions of mitophagy. EMBO J 40, e104705 (2021).

12. Spuch, C., Ortolano, S. & Navarro, C. New insights in the amyloid-Beta interaction with mitochondria. J Aging Res 2012, 324968 (2012).

13. Fang, E.F. et al. Mitophagy inhibits amyloid-beta and tau pathology and reverses cognitive deficits in models of Alzheimer’s disease. Nat Neurosci 22, 401–412 (2019).

14. Palikaras, K., Lionaki, E. & Tavernarakis, N. Coordination of mitophagy and mitochondrial biogenesis during ageing in C. elegans. Nature 521, 525–528 (2015).

15. Pan, J.P. et al. Reduced ULK1 links impaired autophagy and mitophagy to Alzheimer’s disease pathology. Nat Aging 6, 1079–1102 (2026).

16. Markaki, M., Tsagkari, D. & Tavernarakis, N. Mitophagy and long-term neuronal homeostasis. J Cell Sci 136 (2023).

17. McWilliams, T.G. et al. Basal Mitophagy Occurs Independently of PINK1 in Mouse Tissues of High Metabolic Demand. Cell Metab 27, 439–449 e435 (2018).

18. Lee, J.J. et al. Basal mitophagy is widespread in Drosophila but minimally affected by loss of Pink1 or parkin. J Cell Biol 217, 1613–1622 (2018).

19. Koentjoro, B., Park, J.S. & Sue, C.M. Nix restores mitophagy and mitochondrial function to protect against PINK1/Parkin-related Parkinson’s disease. Sci Rep 7, 44373 (2017).

20. Ordureau, A. et al. Temporal proteomics during neurogenesis reveals large-scale proteome and organelle remodeling via selective autophagy. Mol Cell 81, 5082–5098 e5011 (2021).

21. Diwan, A. et al. Unrestrained erythroblast development in Nix-/- mice reveals a mechanism for apoptotic modulation of erythropoiesis. Proc Natl Acad Sci U S A 104, 6794–6799 (2007).

22. Esteban-Martinez, L. & Boya, P. BNIP3L/NIX-dependent mitophagy regulates cell differentiation via metabolic reprogramming. Autophagy 14, 915–917 (2018).

23. Palikaras, K., Lionaki, E. & Tavernarakis, N. Coupling mitogenesis and mitophagy for longevity. Autophagy 11, 1428–1430 (2015).

24. Schmid, E.T., Pyo, J.H. & Walker, D.W. Neuronal induction of BNIP3-mediated mitophagy slows systemic aging in Drosophila. Nat Aging 2, 494–507 (2022).

25. Choi, G.E. et al. BNIP3L/NIX-mediated mitophagy protects against glucocorticoid-induced synapse defects. Nat Commun 12, 487 (2021).

26. Martin-Maestro, P., Gargini, R., Garcia, E., Perry, G., Avila, J. & Garcia-Escudero, V. Slower Dynamics and Aged Mitochondria in Sporadic Alzheimer’s Disease. Oxid Med Cell Longev 2017, 9302761 (2017).

27. Wang, E. et al. Multiscale proteomic modeling reveals protein networks driving Alzheimer’s disease pathogenesis. Cell 188, 6186–6204 e6113 (2025).

28. Mauthe, M. et al. Chloroquine inhibits autophagic flux by decreasing autophagosome-lysosome fusion. Autophagy 14, 1435–1455 (2018).

29. Klionsky, D.J. et al. Guidelines for the use and interpretation of assays for monitoring autophagy (4th edition)(1). Autophagy 17, 1–382 (2021).

30. Um, J.H., Kim, Y.Y., Finkel, T. & Yun, J. Sensitive Measurement of Mitophagy by Flow Cytometry Using the pH-dependent Fluorescent Reporter mt-Keima. J Vis Exp (2018).

31. Tsakiri, E. et al. Tau protein as a regulator of mitochondrial function and dynamics. Proc Natl Acad Sci U S A 123, e2521642123 (2026).

32. Mossink, B. et al. Human neuronal networks on micro-electrode arrays are a highly robust tool to study disease-specific genotype-phenotype correlations in vitro. Stem Cell Reports 16, 2182–2196 (2021).

33. Davis, A.N. & Tanis, J.E. Measuring Caenorhabditis elegans Sensitivity to the Acetylcholine Receptor Agonist Levamisole. J Vis Exp (2022).

34. Mahoney, T.R., Luo, S. & Nonet, M.L. Analysis of synaptic transmission in Caenorhabditis elegans using an aldicarb-sensitivity assay. Nat Protoc 1, 1772–1777 (2006).

35. Cavallini, A. et al. An unbiased approach to identifying tau kinases that phosphorylate tau at sites associated with Alzheimer disease. J Biol Chem 288, 23331–23347 (2013).

36. Dolan, P.J. & Johnson, G.V. The role of tau kinases in Alzheimer’s disease. Curr Opin Drug Discov Devel 13, 595–603 (2010).

37. Van Acker, Z.P., Bretou, M. & Annaert, W. Endo-lysosomal dysregulations and late-onset Alzheimer’s disease: impact of genetic risk factors. Mol Neurodegener 14, 20 (2019).

