## Supplementary figures, tables and online methods for "BNIP3L/NIX-mediated mitophagy limits Alzheimer’s disease pathology and preserves neuronal function"

### 1 Supplementary Figures

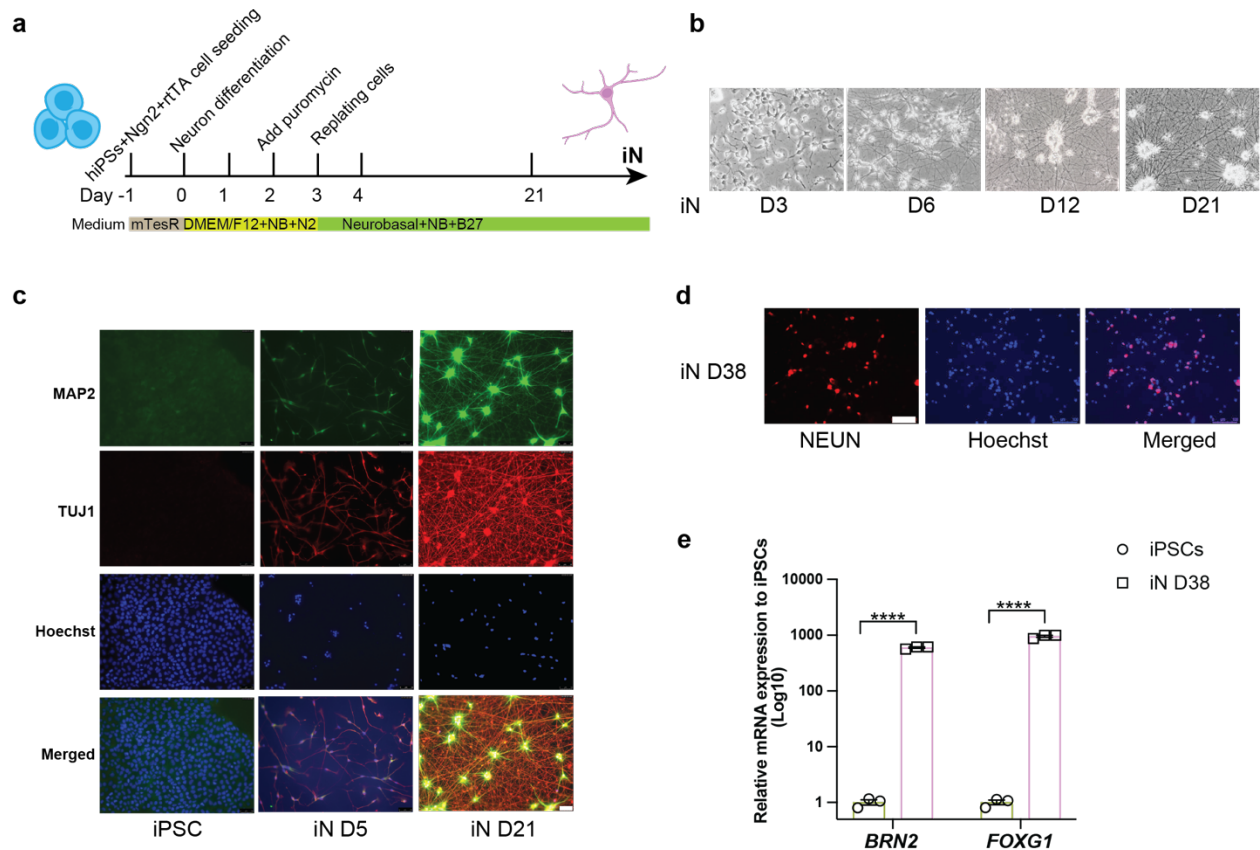

#### 3 Figure S1 (Related to Figure 1)

a. Schematic illustration of neuronal differentiation from iPSCs to iPSC-derived neurons. NB, NT-3 and BDNF.
b. Representative bright-field image of iPSCs cultures at day (D) 3-6-12-21 showing morphological changes during neuronal differentiation.
c. Neuron marker (MAP2, TUJ1) characterization by immunofluorescence in iPSCs, iPSCs-differentiated neuron (iN) at day (D) 5 and day 21. Scale bar, 50 mm.
d. Neuronal marker NEUN expression in iPSC-derived neurons at day 38 detected by immunofluorescence. Scale bar, 100 μm.
e. The relative mRNA expression of excitatory cortical neuron marker *BRN2* and *FOXG1*'s in iPSCs-derived neurons (iN) at differentiation day38 compared to iPSCs.

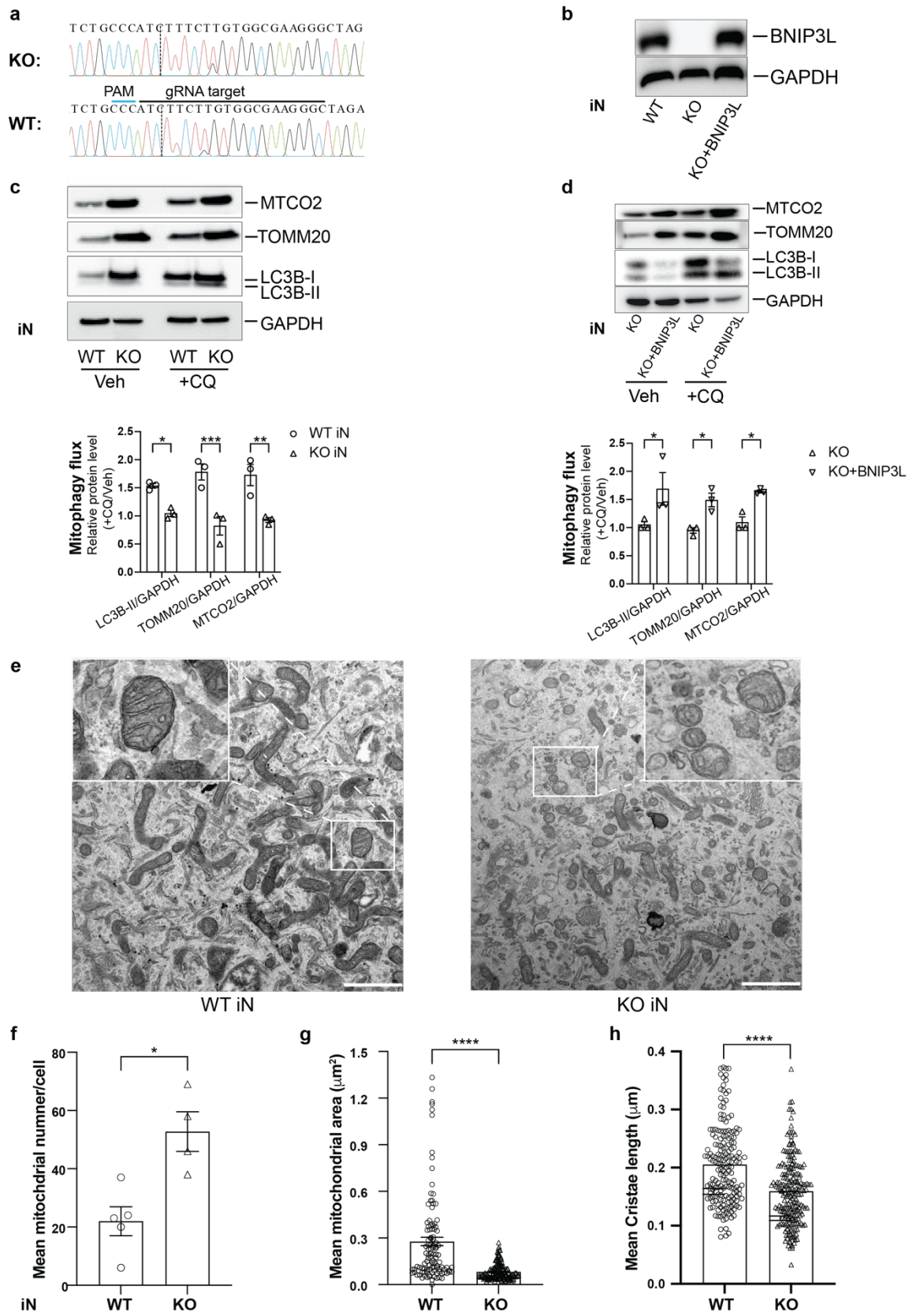

**a.** Sequencing results of Isogenic wild-type control (WT) and BNIP3L knockout (KO) iPSCs generated using CRISPR/Cas9.

**b.** Representative western blot images showing BNIP3L protein expression in iPSC-derived neurons (iN).

**c.** Upper panel showing representative western blot image of LC3B, TOMM20, and MTCO2 expression in WT and KO iN treated with or without 40 μM chloroquine (CQ) for 2 h. Bottom panel showing quantification of mitophagy flux from three batches of independent the Western blot experiments data. Data are presented as mean ± SEM. Statistical significance was determined using two-way ANOVA Šídák's multiple comparisons test. \*\*\* $p < 0.001$.

**d.** Upper panel showing representative western blot image of LC3B, TOMM20, MTCO2, and BNIP3L expression in KO and BNIP3L-overexpressing KO (KO+BNIP3L) iN treated with or without 40 μM CQ for 2 h. Bottom panel showing quantification of the Western blot data from three batches of independent experiments. Data are presented as mean ± SEM. Statistical significance was determined using two-way ANOVA Šídák's multiple comparisons test. \* $p < 0.05$, ns, no significant difference.

**e.** Representative transmission electron microscopy (TEM) images of WT and BNIP3L KO iN. Scale bar, 1 μm. White rectangle showing the magnified image of mitochondria.

**f–h.** Quantification of mean mitochondrial number (f), mean mitochondrial area (g), and mean cristae length (h) in WT and KO iN. Data are presented as mean ± SEM. Statistical significance was assessed using an unpaired *t*-test. \*$P < 0.05$; \*\*\*$P < 0.001$.

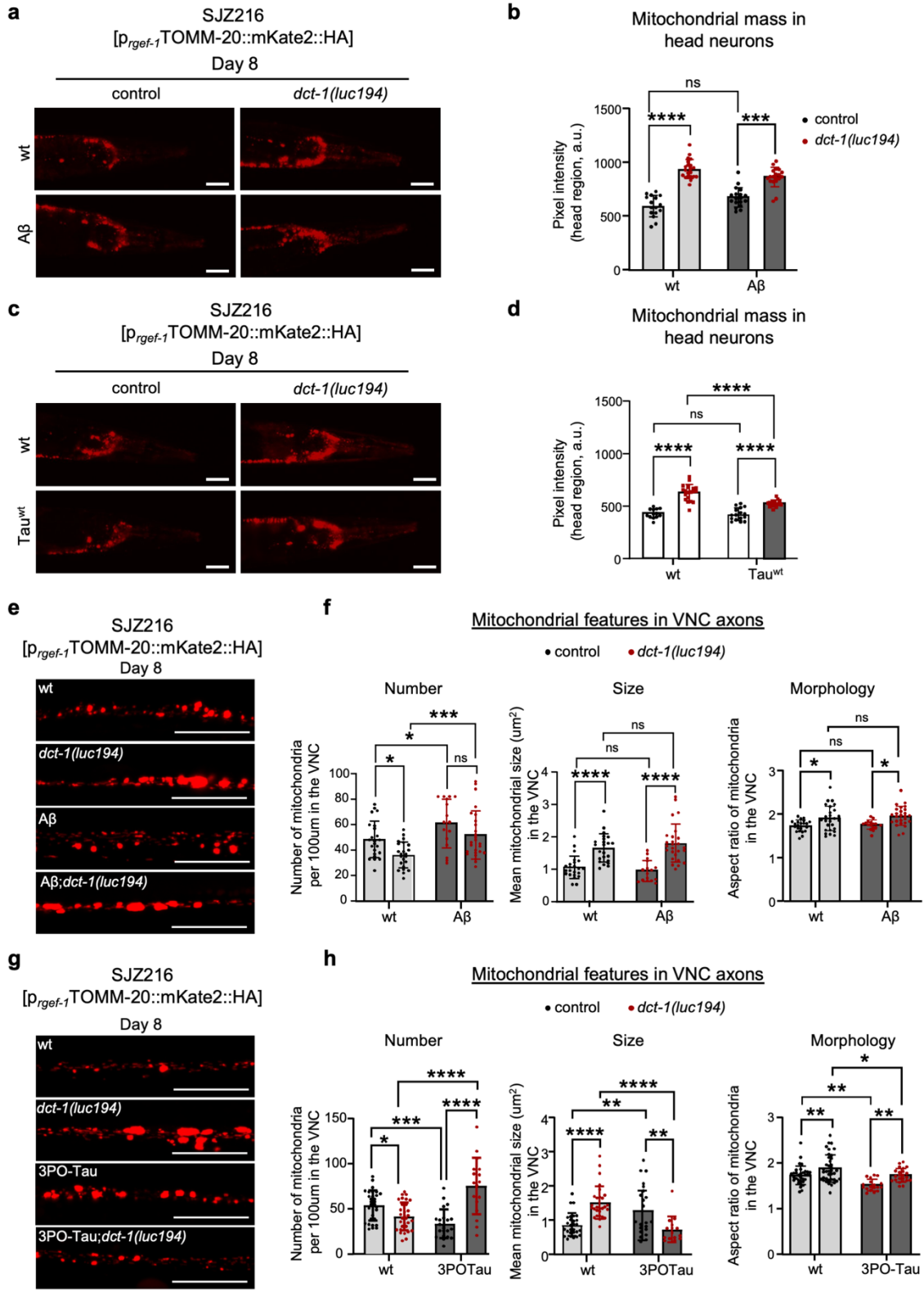

Figure S3 (Related to Figure 3)

a. Representative fluorescence images of the head region of 8-day-old adult nematodes, expressing the pan-neuronal mitochondrial reporter $p_{rgef-1}$ TOMM-20::mKate2::HA (SJZ216 strain) in wild-type and $A\beta$-expressing (GRU102) backgrounds under control or *dct-1(luc194)* mutant conditions. Scale bar, 50 μm.
b. Quantification of neuronal mitochondrial content based on the fluorescence intensity of the pan-neuronal mitochondrial reporter ($p_{rgef-1}$:TOMM-20::mKate2::HA) in the head region of 8-day-old adult *C. elegans* (data presented as mean ± SD; ns $p > 0.05$, \*\*\* $p < 0.001$, \*\*\*\* $p < 0.0001$; one-way ANOVA).
c. Representative fluorescence images of the head region of 8-day-old adult nematodes, expressing the pan-neuronal mitochondrial reporter $p_{rgef-1}$ TOMM-20::mKate2::HA (SJZ216 strain) in wild-type and Tau<sup>wt</sup>-expressing (GRU102) backgrounds under control or *dct-1(luc194)* mutant conditions. Scale bar, 50 μm.
d. Quantification of neuronal mitochondrial content based on the fluorescence intensity of the pan-neuronal mitochondrial reporter ($p_{rgef-1}$:TOMM-20::mKate2::HA) in the head region of 8-day-old adult *C. elegans* (data presented as mean ± SD; ns $p > 0.05$, \*\*\* $p < 0.001$, \*\*\*\* $p < 0.0001$; one-way ANOVA).
e. Representative fluorescence images of axonal mitochondria in 8-day-old adult *C. elegans* expressing the pan-neuronal mitochondrial reporter $p_{rgef-1}$ TOMM-20::mKate2::HA (SJZ216 strain) in wild-type and $A\beta$-expressing (GRU102) backgrounds under control or *dct-1(luc194)* mutant conditions. Scale bar, 20 μm.
f. Quantification of axonal mitochondrial abundance (number of mitochondria per 100um axon length), mitochondrial size and morphology (aspect ratio) in the indicated genotypes (data presented as mean ± SD; ns $p > 0.05$, \* $p < 0.05$, \*\*\*\* $p < 0.0001$; one-way ANOVA).
g. Representative fluorescence images of axonal mitochondria in 8-day-old adult *C. elegans* expressing the pan-neuronal mitochondrial reporter $p_{rgef-1}$ TOMM-20::mKate2::HA (SJZ216 strain) in wild-type and 3POTau-expressing (EVL1372) backgrounds under control or *dct-1(luc194)* mutant conditions. Scale bar, 20 μm.
h. Quantification of axonal mitochondrial abundance (number of mitochondria per 100um axon length), mitochondrial size and morphology (aspect ratio) in the indicated genotypes (data presented as mean ± SD; \* $p < 0.05$, \*\* $p < 0.01$, \*\*\* $p < 0.001$, \*\*\*\* $p < 0.0001$; one-way ANOVA).

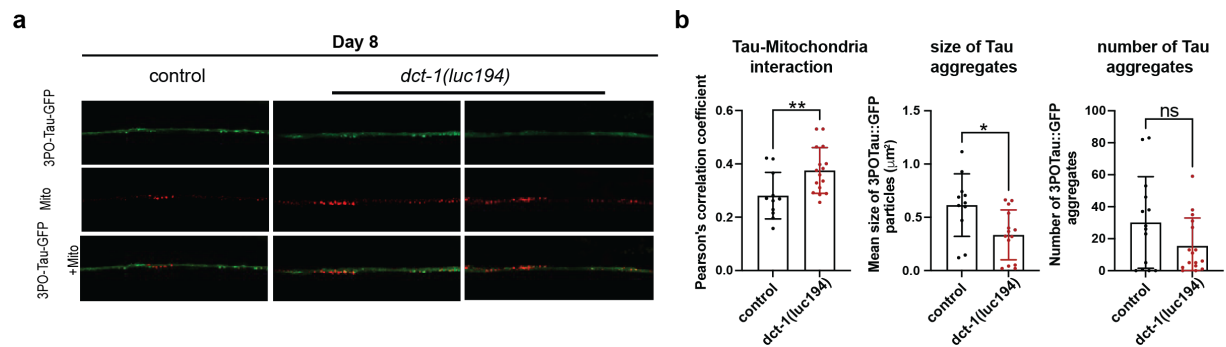

**Figure S4 (Related to Figure 4)**

a. Representative images of the axonal region of 8-day adult nematodes co-expressing the 3POTau::GFP and the TOMM-20::mKate2::HA transgenes (EVL1372;SJZ216 strain) under control or *dct-1(luc194)* mutant conditions.
b. Quantification of the Pearson's correlation coefficient (PCC), number of Tau aggregates and size of Tau aggregates between 3POTau::GFP and the mitochondrial marker TOMM-20::mKate2::HA in the head region of 8-day-old adult nematodes under control or *dct-1(luc194)* mutant conditions. PCC was

calculated to assess the degree of Tau–mitochondria colocalization (data presented as mean ± SD; \*\*
$p < 0.01$; Unpaired t-test).

**a**

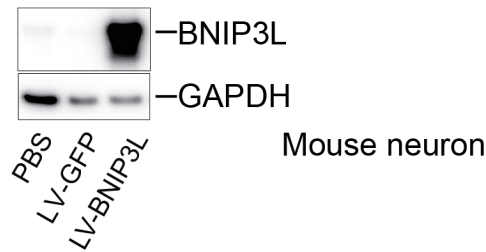

**Figure S5 (Related to Figure 5)**

**a.** Representative Western blot image showing overexpression of human BNIP3L in primary cortical neurons from rTg4510 mice transduced with LV-BNIP3L.

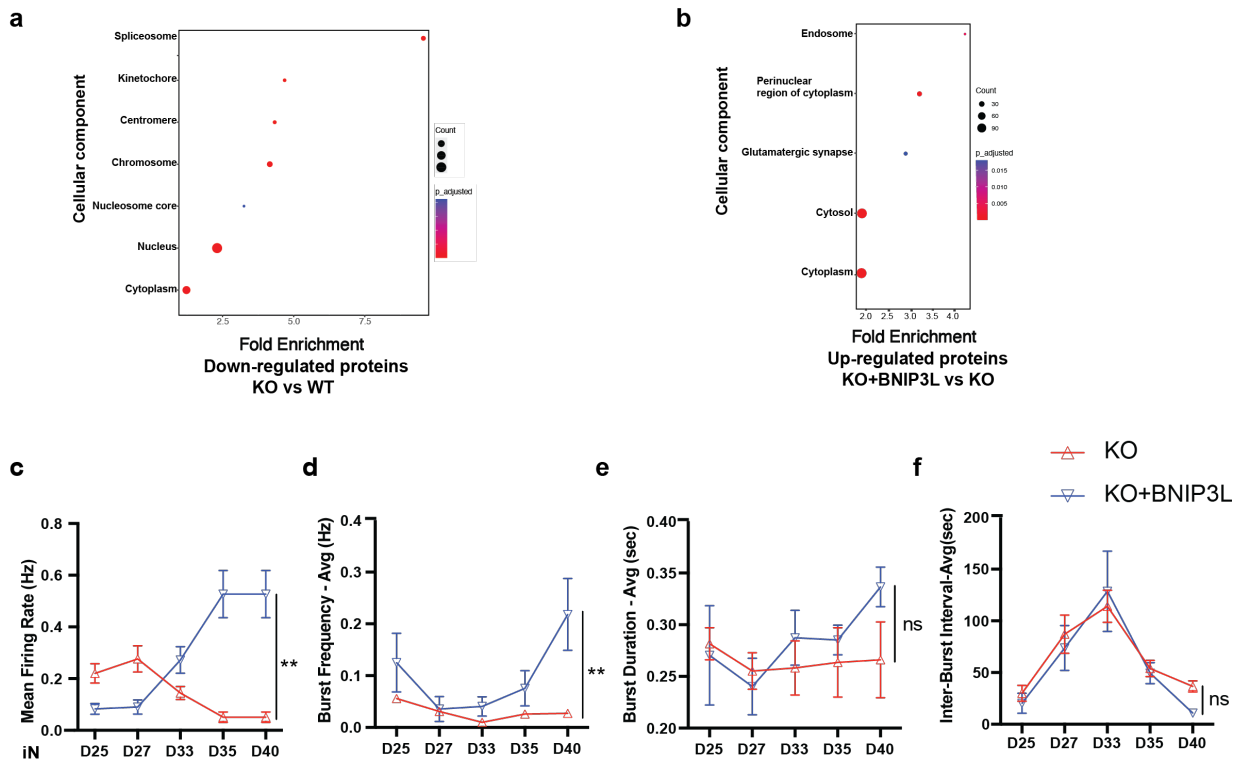

**Figure S6 (Related to Figure 6)**

**a-b.** The gene ontology (GO) cellular component analysis shows the changes associated with down-regulated proteins (a) between KO iN and WT iN and up-regulated proteins (b) between KO iN overexpressed with BNIP3L iN and KO iN. The GO biological processes were organized by Fold enrichment and p<sub>adjusted</sub>.

c-f. Single electrode level MEA data showing (c) mean firing rate (spike rate, Hz). (d) burst frequency-Average (Hz), (e) burst duration-Average (sec) and (f) inter-burst interval-Average (sec) during neuron differentiation time course. Data are presented as mean ± SEM. Statistical analyses were conducted with mixed-effects model with Tukey's multiple comparisons over time with asterisks showing the differences between KO and KO+BNIP3L (KO overexpressed with BNIP3L) iN. \*\* $p < 0.01$, ns, no significant difference.

#### Supplementary tables

##### Supplementary Table 1. *C. elegans* strains used in this study

| Strain name | Parental strains | Genotype | Source |
| --- | --- | --- | --- |
| N2 | - | wild-type Bristol isolate | CGC |
| MLC2543 | - | *dct-1(luc194)* X |  |
| GRU101 | - | *gnals1*[*p<sub>myo-2</sub>*YFP] |  |
| GRU102 | - | *gnals2*[*p<sub>myo-2</sub>*YFP + *p<sub>unc-119</sub>*Abeta1-42] | CGC |
| EVL1654 | - | [*p<sub>rgef-1</sub>*hTau40::GFP] | Brian D. Ackley Lab |
| EVL1372 | - | [*p<sub>rgef-1</sub>*3POTau::GFP + *p<sub>myo-2</sub>*RFP] | Brian D. Ackley Lab |
| KPA426 | GRU101;IR1864 | *gnals1*[*p<sub>myo-2</sub>*YFP]; N2; Ex001[*p<sub>unc-119</sub>*TOMM-20::Rosella + *rol-6(su1006)*] | Palikaras Lab |
| KPA350 | GRU102;IR1864 | *gnals2*[*p<sub>myo-2</sub>*YFP + *p<sub>unc-119</sub>*Abeta1-42]; N2; Ex001[*p<sub>unc-119</sub>*TOMM-20::Rosella + *rol-6(su1006)*] | Palikaras Lab |
| KPA598 | KPA426;MLC2543 | *gnals1*[*p<sub>myo-2</sub>*YFP]; N2; Ex001[*p<sub>unc-119</sub>*TOMM-20::Rosella + *rol-6(su1006)*];*dct-1(luc194)* X | Palikaras Lab |
| KPA597 | KPA350;MLC2543 | *gnals2*[*p<sub>myo-2</sub>*YFP + *p<sub>unc-119</sub>*Abeta1-42]; N2; Ex001[*p<sub>unc-119</sub>*TOMM-20::Rosella + *rol-6(su1006)*];*dct-1(luc194)* X | Palikaras Lab |
| SJZ216 | - | *foxSi44* [*p<sub>rgef-1</sub>*TOMM-20::mKate2::HA::tbb-2 3'UTR] I | CGC |
| KPA346 | SJZ216;MLC2543 | *foxSi44* [*p<sub>rgef-1</sub>*TOMM-20::mKate2::HA::tbb-2 3'UTR] I ; *dct-1(luc194)* X | Palikaras Lab |
| KPA479 | SJZ216;GRU102 | *foxSi44* [*p<sub>rgef-1</sub>*TOMM-20::mKate2::HA::tbb-2 3'UTR] I ; *gnals2*[*p<sub>myo-2</sub>*YFP + *p<sub>unc-119</sub>*Abeta1-42] | Palikaras Lab |
| KPA439 | SJZ216;GRU102;MLC2543 | *foxSi44* [*p<sub>rgef-1</sub>*TOMM-20::mKate2::HA::tbb-2 3'UTR] I ; *gnals2*[*p<sub>myo-2</sub>*YFP + *p<sub>unc-119</sub>*Abeta1-42]; *dct-1(luc194)* X | Palikaras Lab |
| KPA463 | SJZ216;EVL1654 | foxSi44 [*p<sub>rgef-1</sub>*TOMM-20::mKate2::HA::tbb-2 3'UTR] I; [*p<sub>rgef-1</sub>*:hTau40::GFP] | Palikaras Lab |
| KPA500 | SJZ216;EVL1654;MLC2543 | foxSi44 [*p<sub>rgef-1</sub>*TOMM-20::mKate2::HA::tbb-2 3'UTR] I; [*p<sub>rgef-1</sub>*:hTau40::GFP];*dct-1(luc194)* X | Palikaras Lab |
| KPA478 | SJZ216;EVL1372 | foxSi44 [*p<sub>rgef-1</sub>*TOMM-20::mKate2::HA::tbb-2 3'UTR] I; [*p<sub>rgef-1</sub>*13POTau::GFP + *p<sub>myo-2</sub>*RFP] | Palikaras Lab |
| KPA438 | SJZ216;EVL1372;MLC2543 | foxSi44 [*p<sub>rgef-1</sub>*TOMM-20::mKate2::HA::tbb-2 3'UTR] I; [*p<sub>rgef-1</sub>*13POTau::GFP + *p<sub>myo-2</sub>*RFP];*dct-1(luc194)* X | Palikaras Lab |
| KPA445 | EVL1372;MLC2543 | [*p<sub>rgef-1</sub>*13POTau::GFP + *p<sub>myo-2</sub>*RFP];*dct-1(luc194)* | Palikaras Lab |
| KPA446 | GRU102;MLC2543 | *gnals2*[*p<sub>myo-2</sub>*YFP + *p<sub>unc-119</sub>*Abeta1-42]; *dct-1(luc194)* | Palikaras Lab |
| KPA477 |  | N2;Ex[*p<sub>rgef-1</sub>*DCT-1::dct-1 3'UTR + *rol-6(su1006)*] | Palikaras Lab |
| KPA502 | KPA477;MLC2543 | N2;Ex[*p<sub>rgef-1</sub>*DCT-1::dct-1 3'UTR + *rol-6(su1006)*]; *dct-1(luc194)* X | Palikaras Lab |
| JKM2 | - | [*p<sub>rgef-1</sub>*Signalpeptide-Aβ<sub>1-42</sub>::hsp-3(IRES)::wrnScarlet-Aβ<sub>1-42</sub>::*unc-54*(3'UTR)+Prps-0 HydroR] | Kirstein Lab |
| KPA501 | JKM2;MLC2543 | [*p<sub>rgef-1</sub>*Signalpeptide-Aβ<sub>1-42</sub>::hsp-3(IRES)::wrnScarlet-Aβ<sub>1-42</sub>::*unc-54*(3'UTR)+Prps-0 HydroR]; *dct-1(luc194)* X | Palikaras Lab |

98

99

**Supplementary Table 2 Information of iPSCs lines**

|  | Line | Age/sex | Clinical diagnosis |
| --- | --- | --- | --- |
| Ctrl1 | BPC404 | 74/female | Non-affected |
| AD1 | BPC939 | 72/female | Moderate Dementia |
| Ctrl2 | BPC403 | 83/male | Non-affected |
| AD2 | BPC943 | 82/male | Moderate Dementia |

**Supplementary Table 3 qPCR Primers used in this study**

|  | Sequence (5' → 3') | Tm |
| --- | --- | --- |
| *FOXG1-F* | GAGCGACGACGTGTTCATC | 60.9 |
| *FOXG1-R* | GCCGTTGTAACTCAAAGTGCTG | 62.2 |
| *BRN2-F* | CGGCGGATCAAACTGGGATTT | 63 |
| *BRN2-R* | TTGCGCTGCGATCTTGTCTAT | 62.2 |

#### Online Methods

##### iPSCs information and culture condition

iPSCs were obtained with Material Transfer Agreement (MTA) from Zameel Cader group at Nuffield department of Clinical Neurosciences, University of Oxford and from Kristine Freude group at University of Copenhagen (see Supplementary Table 2 for details about age and sex of the patient). iPSCs were cultured in Matrigel (#354277, Fisher Scientific)-coated plates and cultured with mTesR Plus (#100-0276, Stemcell Technologies).

##### iPSCs-derived neurons

iPSCs were differentiated into neurons following a previously validated and published protocol (Zhang et al., 2013). Briefly, iPSCs were cultured in mTesR Plus medium added with Ngn2 and rtTA lentivirus on day -1, and then the next day (Day 0) the culture medium was replaced with DMEM/F12 medium (#31331093, Thermo Fisher) containing N2 (#17502-048, Thermo Fisher), NEAA (#11350912, Fisher Scientific), human BDNF (10 mg/l, #AF-450-02, PeproTech), and human NT-3 (10 mg/l, AF-450-03-1, PeproTech). Doxycycline (2 g/l, #D3072, Clontech) was added on day 0 to induce TetO gene expression. On day 2, a 24 hr puromycin (1 mg/l) selection period was started. On day 3, medium was changed to Neurobasal medium containing B27 (#17504-44, Thermo Fisher), Glutamax (#35050061, Invitrogen), BDNF and NT3; After day 7, 50% of the medium in each well was exchanged every 2 days. Doxycycline was added into the differentiation media during the whole process of neuron differentiation.

#### **Antibodies**

The primary antibodies used were: AMBRA1 (#24907, CST), MUL1 (#NBP1-59068, Novus Biologicals, CO, USA), BNIP3L/NIX (#12396, Cell Signaling Technology, MA, USA), PINK1 (#ab23707, Abcam, MA, USA), LC3B (#NB600-1384SS, Novus Biologicals), GAPDH (#G8795, Sigma, MA, USA), ACTIN (#MA5-11869, Thermo Fisher Scientific), TOMM20 (#WH0009804M1, Sigma), MTCO2 (#A-6404, Thermo Fisher Scientific), TAU (#A0024, DAKO), Phospho-Tau (Ser202, Thr205) Monoclonal Antibody (AT8) (#MN1020, Thermo Fisher Scientific), Phospho-Tau (Thr181) Monoclonal Antibody (AT270) (#MN1050, Thermo Fisher Scientific), Phospho-Tau (Ser356) Polyclonal Antibody (#44-751G, Thermo Fisher Scientific), Phospho-Tau (Ser262) Polyclonal Antibody (#44-750G, Thermo Fisher Scientific).

#### **Western blot**

The cells were lysed in RIPA lysis buffer (#89900, Thermo Fisher Scientific, MA, USA) supplemented with protease inhibitor cocktail (#11836145001, Roche, Basel, Switzerland) and PhosSTOP (#4906837001, Roche) and incubated on ice for 15 min. The protein concentration of the extracts was measured using the BCA protein quantification kit (#23225, Thermo Fisher Scientific). Equal amounts of total protein were loaded onto and resolved by electrophoresis through a Bis-Tris Protein Gel (#NW04125BOX, Thermo Fisher Scientific) and transferred to PVDF membrane (#1620177, Bio-Rad). After blocking with 5% non-fat milk in TBST (50 mM Tris-HCl pH 8.0, 150 mM NaCl, 0.1% Tween 20), the membrane was incubated with 5% non-fat milk containing primary or secondary antibodies at appropriate dilution. The membranes were developed, and the protein signal was detected using the Clarity Max™ ECL Western Blot Detection Kit (#1705062, Bio-Rad). Signals were quantified by densitometry. Constitutively expressed “housekeeping” gene products were used to normalize the data. Image analysis and quantification using Image J software.

#### **Data analysis**

Bar plot in Figure 1C were generated was generated in R studio by analyzing the data from Table S2 page 2 of Wang. *et. al.* Cell, 2025<sup>1</sup>. The association between BNIP3L’s protein expression and AD severity score parameters including Braak stage, Clinical Dementia Rating (CDR), plaque burden, and CERAD scores were analyzed by in-house written R-scripts in R studio.

#### **Lentivirus packaging and infection**

pCHAC-mt-mtKeima (#72342, Addgene), pTet-O-Ngn2-puro (#52047, Addgene), pLV\_hEF1a\_rtTA3 (#62472, Addgene), pMD2.G (#12259, Addgene), pMDLg/pRRE (#12251, Addgene) and pRSV-Rev (#12253, Addgene) were purchased from Addgene. Lentiviruses were packaged in HEK293T cells together with packaging vectors (pCHAC-mt-mKeima (9 µg), pMDLg/pRRE (4.5 µg), pRSV-Rev (4.5 µg), and pMD2.G (4.5 µg)) by transfecting on HEK293 cells using 45 µg 1 ug/uL PEI (polyethylenimine, transfection reagent). Lentivirus-containing media were collected 48 h after transfection and filtered through a 0.45 µm filter to remove the cell debris. The lentiviral preparation was transduced into iPSCs for 8 h. Cells were then grown in normal iPSCs culture medium. hBNIP3L lentivirus (Cat. No. #LVS(VB900139-2286xjg) and EGFP (VB010000-9298rtf) lentivirus was purchased from Vector Builder.

#### **Flow cytometry**

Mitochondrial ROS, mitochondrial mass, and membrane potential in iPSC-derived neurons were assessed using MitoSOX Red (#M36008), MitoTracker Green (#M7514), and TMRM (#T668) (Thermo Fisher Scientific), respectively, following the manufacturer’s instructions. Cells were incubated with 2.5 µM MitoSOX Red, 200 nM MitoTracker Green, or 100 nM TMRM respectively for 15 min at 37 °C in DPBS within a humidified 5% CO<sub>2</sub> incubator. Then cells were washed, harvested, and analyzed using a CytoFLEX flow cytometer (Beckman Coulter). MitoSOX, MitoTracker Green, and TMRM fluorescence were detected using 561/586 nm,

490/516 nm, and standard PE-channel settings, respectively. At least three independent experiments were performed for each assay.

#### **Transmission electron microscopy (TEM)**

Cells were grown to 70% confluence on Thermanox coverslips and fixed with 2% v/v glutaraldehyde in 50 mM sodium phosphate buffer (pH 7.2). The samples were rinsed three times in 0.15 M phosphate buffer (pH 7.2) and post-fixed in 1% w/v OsO<sub>4</sub> in 0.12 M sodium phosphate buffer (pH 7.2) for 2 h. The samples were dehydrated using a graded series of ethanol:water washes, transferred to propylene oxide, and embedded in Epon according to standard procedures. Following polymerization, the Thermanox coverslip was peeled off. Sections, ~60 nm thick, were cut using a Leica UC7 microtome (Leica Microsystems, Vienna, Austria) and collected on copper grids with Formvar supporting membranes, stained with uranyl acetate and lead citrate, and subsequently examined using a Philips CM 100 Transmission EM (Philips, Eindhoven, The Netherlands) operated at an accelerating voltage of 80 kV. Digital images were recorded with slow scan mode on an OSIS Veleta 2k x 2k CCD camera and processed using the ITEM software package. Quantification of mitochondrial number, area and cristae length was done using Image J. The average of mitochondria number was obtained by counting at least 4 different cells and 6 images under different conditions individually. Randomly at least 6 images were used for mitochondrial area quantification. The average of mitochondria cristae length per cell was obtained by counting at least 9 images under different conditions individually.

#### **Mitochondrial bioenergetics**

The Seahorse XFe96 extracellular flux analyzer (Agilent Technologies, CA, USA) was used to measure Oxygen Consumption Rate (OCR) in iPSCs-differentiated neuron. Briefly, neurons were incubated for 1 h in XF media (Agilent Technologies, #103334-100) containing 1 mM sodium pyruvate, 10 mM glucose and 11 mM glutamate (PH7.4, assay medium) at 37°C in a CO<sub>2</sub>-free incubator. Oxygen consumption rates (OCR) were measured at 37 °C using the XFe96 Analyzer from Seahorse Bioscience, which employs extracellular flux analysis in a 96-well plate format. Prior studies have indicated that the presence of oligomycin may lead to an underestimation of maximal mitochondrial respiration<sup>2,3</sup>. To ensure accurate interpretation of the results, two distinct experimental approaches were employed. In the first approach, basal OCR was initially measured, followed by the addition of oligomycin (1.5 μM, #495455, Sigma-Aldrich) to assess respiration-driven ATP synthesis and proton leak-linked OCR. In the second approach, the maximal respiratory capacity was determined through sequential additions of 1.5 μM carbonyl cyanide 4-(trifluoromethoxy) phenylhydrazone (FCCP, #C2920, Sigma-Aldrich), without prior addition of oligomycin. Both protocols included the determination of the fraction of non-mitochondrial oxygen consumption following the addition of 1 μM Rotenone and antimycin A. This ensured a comprehensive assessment of mitochondrial function while avoiding potential misinterpretations of the data.

##### **Quantitative real-time PCR (RT-qPCR)**

RNA was extracted from iPSCs-derived neuron using RNeasy mini kit (#79306, Qiagen, Germany), according to the manufacturer's instructions. RNA purity and concentration were determined by spectroscopy (Agilent BioTek spectrometer, Agilent Technologies). Samples with $OD_{260}/OD_{280} \geq 2.0$ were included in subsequent gene expression analyses. Reverse transcription was performed with the High-capacity cDNA Reverse Transcription Kit (#4368814, ThermoFisher) using 2 μg RNA and random primers. RT-PCR was performed using HOT FIREPol EvaGreen qPCR mix (#08-23-00001, Solis BioDyne) and the QuantStudio 6Flex Real-Time PCR system (Applied Biosystems). Relative mRNA expression was calculated using the $2^{-DDCT}$ method. Primer sequences are shown in Supplementary Table 3.

##### **Generation of BNIP3L knockout iPSCs cell line**

Human induced pluripotent stem cells (hiPSCs, BPC404) were maintained in mTeSR Plus medium (STEMCELL Technologies) on growth factor-reduced Matrigel-coated plates. A BNIP3L-targeting sgRNA (5'-GCCCTTCGCCACAAGAAGAT-3') was used based on the previous literature<sup>4</sup> and cloned into a U6GRNA plasmid. Plasmids were prepared using an endotoxin-free plasmid purification kit, and correct sgRNA insertion was confirmed by Sanger sequencing before transfection<sup>5</sup>.

For genome editing, approximately $8 \times 10^5$ hiPSCs were dissociated into single cells using Accutase and nucleofected with 5 μg CAS9PBKS Cas9-GFP plasmid (Addgene#68371) and 5

µg BNIP3L sgRNA plasmid (backbone vector is Addgene#68370) using the P3 Primary Cell 4D-Nucleofector X Kit (Lonza) and program CB-150. Following nucleofection, cells were recovered in mTeSR Plus medium supplemented with 10 µM Y-27632 and cultured on Matrigel-coated plates. GFP-positive cells were enriched by fluorescence-activated cell sorting (Sony MA900) 48 h after nucleofection and expanded for approximately two weeks.

Single-cell-derived clones were generated by limiting dilution into Matrigel-coated 384-well plates at a theoretical density of 0.5 cell per well and cultured in mTeSR Plus medium containing 10 µM Y-27632. Individual colonies were expanded and screened by PCR amplification of the BNIP3L target locus using the primers 5'-AAGTGCAGAACATTTTGGGAGT-3' and 5'-AATTCGTTTAGAGTGACAGGGC-3'. PCR products were subjected to Sanger sequencing, and editing outcomes were analyzed using ICE (Inference of CRISPR Edits). Clones carrying biallelic frameshift mutations were selected for further experiments.

###### **Proteomic Sample Preparation, Mass Spectrometry and Data Analysis**

Proteomic sample preparation was performed as previously described with minor adjustments<sup>6</sup>. Briefly, iPSC-derived neuronal suspensions (2,000 cells in 5 µL) were prepared in technical triplicates for each sample with a digestion master mix buffer containing 0.2% n-dodecyl-β-D-maltoside (DDM), 100 mM triethylammonium bicarbonate (TEAB), 20 ng/µL trypsin, and 10 ng/µL Lys-C. Samples were incubated at 37°C for 3 h, acidified with formic acid to a final concentration of 0.1%, and loaded onto EvoTips according to the manufacturer's protocol. Approximately 20 ng of peptides in 5 µL were loaded onto each EvoTip. Peptides were analyzed using an Evosep One LC system operated with the 80 samples-per-day (SPD) method and coupled to a Thermo Fisher Orbitrap Astral mass spectrometer using data-independent acquisition (DIA). Raw DIA-MS data were analyzed in Spectronaut (Biognosys) for protein identification and quantification. The resulting protein abundance matrix was exported for downstream filtering, statistical analyses, and visualization in R. Protein identifications corresponding to non-human contaminants were removed prior to downstream analyses. Samples were grouped into three biological replicate sets (samples 1–3, 4–6, and 7–9), and proteins with more than one missing value within any replicate set were excluded. Proteins with isolated missing values across different replicate sets were retained. After filtering, 6,823 proteins were identified across all samples. Functional enrichment analyses were performed using the DAVID Gene Ontology platform and custom R scripts. Raw mass spectrometry data have been deposited to the ProteomeXchange Consortium via the PRIDE repository under accession number [accession number].

##### **Confocal microscopy**

Differentiated neurons were grown on the μ-Slide 8-well chamber slide (#80826, ibidi, Germany) stained with mitophagy detection kit (#MD01-10, Dojindo, Japan) in a humidified incubator under 5% CO<sub>2</sub> at 37°C following kit's instructions. After washing, neurobasal media without phenol red (#11570426, Thermo Fisher Scientific) was added for live-cell imaging. Images were acquired using the LSM780 confocal microscope system with a 63x oil immersion objective lens. Mitochondria were labelled using Pk-mito (#SC053, SPIROCHROM) at 2000X dilution of the stock for 15 min. Nuclei were stained with Hoechst 333425 (5 μM) for 10 min at RT. After washing, neurobasal media without phenol red (#11570426, Thermo Fisher Scientific) was added for live-cell imaging. Z-stack images were acquired using the LSM780 confocal microscope system with a 63x oil immersion objective lens.

##### **Microelectrode array (MEA)**

iPSC-derived neurons were plated in CytoView MEA 96-well plates (#1001510, Axion BioSystems). Wells were sequentially coated with 0.1 M polyethylenimine (PEI) in 0.1 M borate buffer for 1 h at 37 °C, washed three times with ultrapure water, air-dried, and subsequently coated with laminin (1 μg ml<sup>-1</sup>) overnight at 37 °C. Neuronal suspensions droplet (10 μl per well; ~10,000 cells) were seeded onto the center of coated well and allowed to attach for 1 h before differentiation medium was added to a final volume of 150 μl per well.

Neuronal network activity was recorded throughout differentiation using the Maestro MEA platform (Axion BioSystems) at a sampling rate of 12.5 kHz under temperature-controlled conditions (37 °C). Spontaneous activity was monitored twice a week by acquiring 10-min recordings for up to 6 weeks. Before each recording session, plates were equilibrated for 3–5 min, and culture medium was refreshed 24 h beforehand<sup>7</sup>.

##### **Neurite outgrowth staining**

Neurite outgrowth and cell viability were assessed in iPSC-derived neurons cultured in 96-well plates (#165305, Thermo Fisher Scientific) using the Neurite Outgrowth Staining Kit (#A15001, Thermo Fisher Scientific) according to the manufacturer's instructions. Following removal of culture medium, neurons were incubated with Cell Viability Indicator and Cell Membrane Stain (1:1,000 dilution each) for 15 min at 37 °C in 5% CO<sub>2</sub>. After staining, cells were washed and incubated with Background Suppression Solution. Fluorescence was quantified using a Synergy H1 plate reader (#17032723, BioTek) with excitation/emission settings of 483/525 nm for the viability indicator and 554/567 nm for the membrane stain.

##### **Aβ42/Aβ40 and Aβ42/Aβ38 measurement**

Conditioned media from iPSC-derived neurons were collected, clarified by centrifugation (1,200 × g, 10 min, 4 °C), aliquoted, and stored at -80 °C until analysis. For each experiment, culture supernatants from three independent experiment were analyzed and each individual sample has 5 replicates. Aβ42/Aβ40 and Aβ42/Aβ38 ratios were quantified using the V-PLEX Plus Aβ Peptide Panel 1 (6E10) Kit (K15200G-1, MSD) according to the manufacturer's instructions. Data were analyzed using MSD Discovery Workbench software.

##### **Preparations of pathological tau seeds from rTg4510 mouse**

The rTg4510 transgenic mice overexpress human 4R0N mutant P301L tau in addition to endogenous murine tau largely restricted to the forebrain by the CaMKIIα promoter<sup>8</sup>. Snap-frozen brains from 32-40 weeks old rTg4510 mice with advanced tau pathology in the forebrain were pooled and forebrain homogenates were separated by differential centrifugation into Tris-buffered saline (TBS)-extractable (S1), S1 precipitate (S1p), and sarkosyl-insoluble pellet (P3) fractions as established previously<sup>9</sup>. Tau seeds from soluble S1p (oligomeric tau species) and insoluble P3 (fibrillar tau species) fractions were prepared as described previously<sup>10</sup>. Briefly, protease and phosphatase inhibitors and ion chelators in the buffers were omitted and forebrain homogenates were separated by centrifugation at 27,000 × g for 20 min at 4°C in S1. The pellet was re-homogenized in 5 volumes of high salt/sucrose buffer and centrifuged as above. The supernatant was collected and incubated with 1% sarkosyl for 1 h at 37°C, followed by centrifugation at 150,000 × g for 1 h at 4°C to obtain P3. S1 was separated by further centrifugation at $150,000 \times g$ for 20 min at $4^{\circ}\text{C}$ to obtain S1p. Tau concentrations in the different fractions were determined with the Innotest human total TAU ELISA (Fujirebio, Tokyo, Japan) following manufacturer's instructions.

##### **Seeded tau aggregation and lentiviral BNIP3L/NIX overexpression in cortical cultures from rTg4510 mice**

Murine cortical neurons (CTX) were isolated from day E14-16 rTg4510 mouse embryos as described previously<sup>10</sup>. The generation of rTg4510 mice and the primer pairs to genotype rTg4510 embryos were described previously<sup>8</sup>. Briefly, single transgenic CaMKIIα promoter tTA activator mice were time-mated with single transgenic human mutant tau responder mice. Pregnant females were euthanized and embryos were genotyped with brain DNA using the primer pair's 5'-AGGCTGCTCTACACCTAGCT-3' and 5'-CAGCGCATTAGAGCTGCTTA-3' for the tTA activator transgene and 5'-CCAACGCCACCAGGATTC-3' and 5'-AGCTGGGTGGTGTCTTTGGA-3' for the human mutant tau transgene while isolated embryonic cortices were kept in Hibernate E without calcium chloride (BrainBits LLC, Springfield, IL, USA) at $4^{\circ}\text{C}$. Cortices from rTg4510 mouse embryos were selected and dissociated neurons plated on 100 $\mu\text{g}/\text{ml}$ poly-L-lysine coated dishes at a density of $0.13 \times 10^6$ cells/cm<sup>2</sup> (400,000 cells/ml, 100 $\mu\text{l}/\text{well}$, 96-well plate) and cultured in Neurobasal plus medium supplemented with 2% B-27 plus with antioxidants, 0.5 mM L-glutamine, 100 U/ml penicillin, and 0.1 mg/ml streptomycin, all solutions from Gibco-BRL Invitrogen (ThermoFisher Scientific, Waltham, MA, USA). At 4 days in vitro (DIV) half of the medium was replaced with fresh Neurobasal plus medium and 1 $\mu\text{M}$ cytosine arabinoside added to halt proliferating cells. Thereafter half of the medium was replaced with fresh medium every third day. The proportion of glia cells in the cultures was less than 10%, as assessed by an antibody against glia-fibrillary-acidic protein.

CTX were transduced at DIV 1 with lentiviral constructs from VectorBuilder Inc. (Chicago, IL, USA) containing human BNIP3L/Nix (vector ID: VB900139-2286xjg) or EGFP (VB010000-9298rtf) at final titers of $4 \times 10^8$, $4 \times 10^7$, $4 \times 10^6$ and $4 \times 10^5$ virus particles/well or incubated with a similar volume PBS directly in the Neurobasal plus medium. Transduction efficiency was estimated by counting green fluorescent protein (GFP) positive cells at DIV7: about 50-90% GFP positive cells were observed at the two highest titers. Final titers of $4 \times 10^8$ virus

particles/well were applied for most tau seeding experiments. At DIV 7, rTg4510 CTX were treated with 0.2 ng S1p or P3 tau seeds or left unseeded incubated with a similar volume PBS (naive). At DIV 8, a complete medium change was performed to remove any residual tau seeds not taken up, at DIV 12 cultures were fed with fresh Neurobasal plus medium and at DIV 15, neurons were harvested to analyze seeded tau aggregation by homogeneous time resolved fluorescence (HTRF) tau aggregation immunoassay and tau immunocytochemistry as described previously<sup>10</sup>.

###### **Measurement of seeded tau aggregation in CTX by HTRF tau aggregation immunoassay and tau immunocytochemistry**

The HTRF human tau aggregation immunoassay kit (#6FTAUPEG, Cisbio, Codolet, France) is based on the principle of time-resolved measurement (TR) of fluorescence resonance energy transfer (FRET) between two fluorophores in proximity coupled to a pair of antibodies recognizing human tau. Using the same monoclonal antibody for the donor and acceptor fluorophores ensures that a TR-FRET signal is only obtained when the epitope is in a multimer form, thus quantifying selectively human tau aggregates. Cultures were lysed per well in 50 μl ice-cold lysis buffer (1% Triton X-100 in 50 mM Tris, 150 mM NaCl (pH 7.6) supplemented with 1% protease inhibitor mixture (Roche, Basel, Switzerland), 1% phosphatase inhibitor cocktail I and II, and 0.2% benzonase under shaking at 200 rpm for 45 minutes at 4°C. Optimal sample dilutions were established according to manufacturer's protocol. In brief, rTg4510 lysates were diluted 4-fold in lysis buffer, and 9 μl sample was mixed with 4.5 μl terbium (Tb)-cryptate donor fluorophore and 4.5 μl d2 acceptor fluorophore coupled human tau antibody. After 20 h incubation, HTRF signal was measured on a PHEARstar (BMG LABTECH, Ortenberg, Germany) using a pulsed 337 nm laser excitation and time delayed simultaneous dual emission at 665/620 nm. The data are presented as percent tau aggregation above background normalized to protein concentrations (μg/μL) measured in the bicinchoninic acid (BCA) Protein Assay Kit from Pierce (ThermoFisher Scientific, Waltham, MA, USA) according to manufacturer's instructions. For detection of cell viability, the percentage of viable cells in the neuronal cultures was quantified at DIV15 before cell lysis with the resazurin assay kit (Abcam, Cambridge, UK) per manufacturer's instructions. Each condition was measured in at least 5 replicates (n = 5 wells) per culture plate. For detection of human BNIP3L/NIX overexpression and total tau, 1 μg total protein from the rTg4510 lysates was used for western blotting as described previously<sup>11</sup>. BNIP3L/NIX was detected using the rabbit polyclonal BNIP3L/NIX (1:1000, #12396, Cell Signaling Technology, Danvers, MA, USA) and tau was detected using the rabbit polyclonal DAKO total tau antibody (1:5000, A0024, Agilent, Santa Clara, CA, USA) and the rabbit polyclonal E1 antibody. The E1 antibody was raised against amino acids 19–33 (GLGDRKDQGGYTMHQ) of the longest isoform of human tau 2N4R<sup>12</sup> and recognizes exclusively human tau. The DAKO tau antibody recognizes both human and murine tau.

For immunocytochemistry, CTX were fixed with ice-cold v/v 80% methanol for 10 min and washed in PBS at DIV15. Blocking was performed in 1% BSA for 1 h, and cultures were subsequently incubated with the mouse tau antibody AT8 (1:500, MN1020, Invitrogen, ThermoFisher Scientific, Waltham, MA, USA) and the rabbit BNIP3L/Nix antibody (1:500, #12396, Cell Signaling Technology, Danvers, MA, USA) overnight at 4°C. After primary antibody staining, cultures were washed in PBS and incubated with secondary anti-mouse (1:1000, A31570, Alexa555-donkey-anti-mouse, Invitrogen, ThermoFisher Scientific, Waltham, MA, USA) and anti-rabbit (1:1000, 715-545-152 Alexa488-donkey-anti-rabbit, Jackson ImmunoResearch Europe Ltd, Ely, UK) antibodies and Hoechst 33342 (1:500, ThermoFisher Scientific, Waltham, MA, USA) for 1 hour at room temperature. After two washing steps in PBS, cultures were subjected to high-content imaging with a Cellomics ArrayScan VTI HCS Reader (ThermoFisher Scientific, Waltham, MA, USA) equipped with a 20x/0.70 objective (Olympus, Tokyo, Japan) and analysed in ThermoScientific HCS Studio: Cellomics Scan Version 6.6.0 (ThermoFisher Scientific, Waltham, MA, USA). Each condition was measured in 5 replicates (5 = wells) per culture plate, with each well imaged in 16 fields and an algorithm (Thermo Scientific HCS Studio: Cellomics Scan Version 6.6.3. Assay template: SpotDetector.v4) was created to analyse number, size, and intensity of AT8-positive inclusions. Seeded tau aggregation was measured as AT8-positive Spot Total Intensity/viable cells. The same algorithm was used to analyse human BNIP3L/Nix expression. Methanol fixation impaired the GFP signal, which analysis was omitted at DIV15. Viable cell count was performed at a separate re-scan only counting Hoechst positive cell bodies from cells that were alive before fixation, based on nuclear size and fluorescent intensity. Each condition was measured in 5 technical replicates (wells) per culture experiment, with each well imaged in 16 fields. Confocal images were acquired with a Leica DMI 8 microscope equipped with a 20x/0.45 objective (HC

PL Fluotar, Leica, Wetzlar, Germany). All data analyses were performed using GraphPad Prism version 9 (GraphPad Software, San Diego, CA, USA). Comparisons involving three groups were analyzed using Brown–Forsythe and Welch ANOVA, followed by Dunnett’s T3 post hoc test, and statistical significance is indicated as $p < 0.0001$ (\*\*\*\*).

##### ***C. elegans* strains and maintenance**

All *C. elegans* strains were maintained on NGM plates seeded with *Escherichia coli* OP50 at 20°C using standard procedures<sup>13</sup>. All *C. elegans* strains used in this study are shown in Supplementary Table 1.

##### **Nematode strain generation**

To generate the plasmid construct for the pan-neuronal expression of DCT-1, the *rgef-1* promoter (approximately 3,5kb upstream of the *rgef-1* gene) was first isolated from N2 genomic DNA using the following primers which include PstI and NheI restriction sites: *rgef-1*\_prom\_FW: 5’-CTGCAGCGTTTCCGATACCCCCTTATATC-3’ and *rgef-1*\_prom\_RV: 5’-GGTACCACGACGAACGATTGAGCAG-3’ and ligated to the PCR-II TOPO 2.1 vector (TOPO™ TA Cloning™ Kit, ThermoFisher Scientific). The *rgef-1* promoter was isolated from the TOPO vector with PstI and NheI sites and was subsequently ligated to the L2528 (pD96.41 vector, Andrew Fire plasmids), already digested with the same enzymes (removal of the *mec-7* promoter). Next, the *dct-1* genomic region, including its endogenous 3’ untranslated region (3’UTR), was isolated from N2 genomic DNA using the following primers: *dct-1*\_NheI\_FW: 5’-GCTAGCATGTCCTCATTTCTTGAGTTTG-3’ and *dct-1*\_AgeI\_RV: 5’-ACCGGTATGCCCAGGTAACAGCAAAG-3’ and ligated to the PCRII-TOPO 2.1 vector. Finally, the *dct-1* genomic region was excised from TOPO vector by digestion with NheI and AgeI and ligated to the “*rgef-1* promoter in pD96.41” vector, already linearized with the same enzymes, leading to the generation of the final “*p<sub>rgef-1</sub>*DCT-1::*dct-1* 3’UTR” plasmid. For the generation of transgenic animals, the resulting “*p<sub>rgef-1</sub>*DCT-1::*dct-1* 3’UTR” construct was microinjected into the gonads of young adult N2 hermaphrodites together with pRF4 [*rol-6(su1006)*] as a co-transformation marker. Transgenic progeny were identified by the dominant roller phenotype and maintained as an extrachromosomal array.

#### Assessment of neuronal mitophagy in *C. elegans*

Neuronal mitophagy was assessed using the mitochondria-targeted Rosella biosensor TOMM-20::Rosella expressed under the pan-neuronal *unc-119* promoter, as previously described<sup>14</sup>. The reporter consists of the pH-sensitive GFP and pH-stable DsRed fluorescent proteins fused to the mitochondrial outer membrane protein TOMM-20, enabling mitophagy to be monitored based on the differential response of the two fluorophores to lysosomal acidification. Mitophagy was analyzed in KPA426 [gnaIs1[pmyo-2::YFP]; Ex001[punc-119::TOMM-20::Rosella + rol-6(su1006)]] control animals and KPA350 [gnaIs2[pmyo-2::YFP + punc-119::Aβ1-42]; Ex001[punc-119::TOMM-20::Rosella + rol-6(su1006)]] Aβ-expressing animals, in the presence or absence of the *dct-1(luc194)* mutation. Fluorescence images were acquired in the GFP and DsRed channels at 10× magnification, and neuronal mitophagy was quantified in the tail region to avoid interference from the strong pharyngeal myo-2::YFP co-injection marker present in the GRU101 and GRU102 backgrounds. Image analysis was performed using Fiji/ImageJ. Following separation of the GFP and DsRed channels, an equivalent region of interest (ROI) encompassing the tail was defined for each animal and applied to both fluorescence channels. Mean GFP and DsRed fluorescence intensities were measured within the corresponding ROI, and neuronal mitophagy was expressed as the GFP:DsRed fluorescence intensity ratio. Because GFP fluorescence is quenched upon delivery of the reporter to the acidic lysosomal compartment while DsRed fluorescence remains relatively stable, lower GFP:DsRed ratios indicate increased mitophagy, whereas higher ratios indicate reduced mitophagy.

#### Assessment of mitochondrial number, size and morphology in the axons of the ventral nerve cord in *C. elegans*

Mitochondrial number, size and morphology within the ventral nerve cord (VNC) axons were quantified by using the *C. elegans* strains: SJZ216, KPA346, KPA479, KPA439, KPA478 and KPA438. For each condition, 20 nematodes at defined adult stages (Day 5/young; Day 8/aged) were immobilized on a slide in a drop of 40mM tetramisole hydrochloride, mounted under a coverslip, and sealed. Confocal imaging of mKate2-labeled mitochondria in the VNC between the head and vulval regions was performed using a Zeiss LSM900 microscope with a 40× objective under identical acquisition settings. All images used for quantitative comparisons were acquired using identical imaging parameters. Two images encompassing distinct VNC regions were acquired per animal. Image analysis was performed using Fiji/ImageJ (NIH). A 100-μm segment of the VNC was defined as the region of interest (ROI), and mKate2-positive mitochondrial profiles were identified and quantified using the Analyze Particles function. Mitochondrial abundance was determined as the number of individual mitochondrial profiles within each 100-μm VNC segment, whereas mitochondrial size was determined from the area of each detected mitochondrial profile. Mitochondrial morphology was further assessed by calculating the aspect ratio (AR) of individual mitochondria, defined as the ratio of the major (Feret) to the minor axis (minFeret). An AR approaching 1 indicates a more rounded mitochondrial profile, whereas increasing AR values indicate progressively elongated mitochondria. Graph generation and statistical analyses were performed using GraphPad Prism 9.0.0 (GraphPad Software, San Diego, CA, USA).

###### **Analysis of Tau aggregates and their colocalization with mitochondria in *C. elegans* ventral nerve cord axons**

The number and size of Tau aggregates in the axons of the ventral nerve cord were assessed as described above, using the strains: KPA478 and KPA438. Images were acquired in the GFP channel, and 3POTau::GFP aggregates were subsequently quantified for number and size. For each condition, 20 nematodes at defined adult stages (Day 5, young; Day 8, aged) were immobilized as previously described and two images were acquired in both green and red channels from the same region of the ventral nerve cord (head to vulva). 3POTau::GFP aggregates were subsequently quantified for number and size using the Particle Analysis function in ImageJ. For the colocalization analysis with mitochondria, the same regions were imaged in both the red and green channels, and composite images were analyzed using the JACoP plugin in ImageJ to calculate Pearson's correlation coefficients<sup>15</sup>. Graphs and statistical analysis were generated using GraphPad Prism 9.0.0 (GraphPad Software, San Diego, California USA, www.graphpad.com).

###### **Quantification of Tau and Aβ aggregation in *C. elegans***

Tau and Aβ aggregation was assessed in the head region of the respective transgenic *C. elegans* models (EVL1372, KPA445, JKM2, KPA501). Young day-5 animals were immobilized in 40mM tetramisole hydrochloride on microscope slides, mounted under coverslips and sealed prior to imaging. Fluorescently labeled 3POTau::GFP- or Aβ::wrmScarlet- positive particles were imaged in the head region using a Zeiss LSM 900 confocal microscope under identical acquisition settings for all genotypes within each experimental comparison. For each animal, the head region was defined as the region of interest (ROI), and fluorescent Tau- or Aβ-positive particles were identified using a consistent intensity threshold applied across the corresponding experimental groups. Following image segmentation, the number and size of fluorescent particles were quantified using the Analyze Particles function in Fiji/ImageJ. The same image-processing and particle-detection parameters were maintained across all genotypes within each experiment. Graph generation and statistical analyses were performed using GraphPad Prism 9.0.0 (GraphPad Software, San Diego, CA, USA).

#### **Olfactory associative learning and memory assay**

Associative learning and memory were assessed using an isoamyl alcohol (IA)-dependent olfactory chemotaxis assay adapted from Cao *et al.* 2023<sup>16</sup>. Age-synchronized animals were generated by hypochlorite synchronization and maintained on NGM plates seeded with *E. coli* OP50 at 20 °C until day 1 of adulthood. Prior to the assay, animals of each genotype were collected in M9 buffer, washed thoroughly and divided equally into naïve and conditioned groups. Animals were transferred to unseeded NGM plates and subjected to 90 min of food deprivation at 20 °C. For associative conditioning, 10 μl of undiluted IA was applied to the inner surface of the plate lid during the starvation period, whereas naïve animals underwent identical food deprivation in the absence of IA. Following conditioning, animals were collected in M9 buffer and transferred to chemotaxis assay plates consisting of 8.5-cm NGM plates without bacterial food. Each assay plate contained a starting position (S), an IA attractant position (IA) and a control position (T). The IA and control positions were each located 4.5 cm from the starting point and 6.5 cm from one another. At least 15 min before the assay, 15 μl of 20 mM sodium azide was applied to both the IA and control positions to immobilize animals reaching either location. Animals were placed at the starting position, and 4 μl of IA diluted 1:50 in water was applied at the attractant position. Plates were immediately sealed and incubated at 20 °C for 2 h. Animals at the IA, control and starting regions were subsequently counted, and the chemotaxis index (CI) was calculated as: $CI = (N - N) / (N + N + N)$, where N, N and N represent the numbers of animals at the IA, control and starting regions, respectively. The assay was performed in three independent biological replicates, with one plate analyzed per genotype and condition in each experiment.

##### **Pharmacological paralysis assays for assessment of neurotransmission in *C. elegans***

Aldicarb and levamisole paralysis assays were performed on NGM plates to assess neuromuscular transmission<sup>17, 18</sup>. Aldicarb inhibits acetylcholinesterase, resulting in progressive accumulation of acetylcholine at the neuromuscular junction; the kinetics of aldicarb-induced paralysis were therefore used as an indirect readout of presynaptic cholinergic transmission. Levamisole-induced paralysis was used as a complementary measure of postsynaptic neuromuscular responsiveness, as levamisole directly activates nicotinic acetylcholine receptors at the neuromuscular junction. Freshly prepared NGM plates were supplemented with 1mM aldicarb or 0,4mM levamisole and allowed to equilibrate at room temperature before use. Plates were subsequently seeded with 50uL *E. coli* OP50. 30-40 age-synchronized animals of the indicated genotypes were transferred onto drug-containing plates, and paralysis was scored at regular time intervals. Animals were considered paralyzed when they failed to move in response to gentle mechanical stimulation. The percentage of paralyzed animals was calculated at each time point relative to the total number of animals analyzed. Paralysis curves were generated and statistically analyzed using GraphPad Prism 9.0.0 (GraphPad Software, San Diego, CA, USA). The assay was performed in at least three independent biological replicates

#### **References**

1. Wang, E. *et al.* Multiscale proteomic modeling reveals protein networks driving Alzheimer's disease pathogenesis. *Cell* **188**, 6186-6204 e6113 (2025).
2. McNally, L.A., Altamimi, T.R., Fulghum, K. & Hill, B.G. Considerations for using isolated cell systems to understand cardiac metabolism and biology. *J Mol Cell Cardiol* **153**, 26-41 (2021).
3. Ruas, J.S., Siqueira-Santos, E.S., Amigo, I., Rodrigues-Silva, E., Kowaltowski, A.J. & Castilho, R.F. Underestimation of the Maximal Capacity of the Mitochondrial Electron Transport System in Oligomycin-Treated Cells. *PLoS One* **11**, e0150967 (2016).
4. Ordureau, A. *et al.* Temporal proteomics during neurogenesis reveals large-scale proteome and organelle remodeling via selective autophagy. *Mol Cell* **81**, 5082-5098 e5011 (2021).

5. Tian, W. *et al.* Cell-based glycoengineering of extracellular vesicles through precise genome editing. *N Biotechnol* **83**, 101-109 (2024).
6. Ye, Z. *et al.* One-Tip enables comprehensive proteome coverage in minimal cells and single zygotes. *Nat Commun* **15**, 2474 (2024).
7. Mossink, B. *et al.* Human neuronal networks on micro-electrode arrays are a highly robust tool to study disease-specific genotype-phenotype correlations in vitro. *Stem Cell Reports* **16**, 2182-2196 (2021).
8. Santacruz, K. *et al.* Tau suppression in a neurodegenerative mouse model improves memory function. *Science* **309**, 476-481 (2005).
9. Sahara, N. *et al.* Characteristics of TBS-extractable hyperphosphorylated tau species: aggregation intermediates in rTg4510 mouse brain. *J Alzheimers Dis* **33**, 249-263 (2013).
10. Volbracht, C. & Fog, K. Deubiquitinating Enzymes Ubiquitin-Specific Proteases 7 and 10 Regulate TAU Aggregation. *Int J Mol Sci* **26** (2025).
11. Rostgaard, N., Jul, P.H., Garmer, M. & Volbracht, C. Increasing O-GlcNAcylation Attenuates tau Hyperphosphorylation and Behavioral Impairment in rTg4510 Tauopathy Mice. *J Integr Neurosci* **22**, 135 (2023).
12. Crowe, A., Ksiazek-Reding, H., Liu, W.K., Dickson, D.W. & Yen, S.H. The N terminal region of human tau is present in Alzheimer's disease protein A68 and is incorporated into paired helical filaments. *Am J Pathol* **139**, 1463-1470 (1991).
13. Stiernagle, T. Maintenance of *C. elegans*. *WormBook*, 1-11 (2006).
14. Palikaras, K., Lionaki, E. & Tavernarakis, N. Mitophagy Dynamics in *Caenorhabditis elegans*, in *Autophagy: Methods and Protocols*. (eds. N. Ktistakis & O. Florey) 655-668 (Springer New York, New York, NY; 2019).
15. Bolte, S. & Cordelières, F.P. A guided tour into subcellular colocalization analysis in light microscopy. *J Microsc* **224**, 213-232 (2006).
16. Cao, S.-q., Wang, H.-L., Palikaras, K., Tavernarakis, N. & Fang, E.F. Chemotaxis assay for evaluation of memory-like behavior in wild-type and Alzheimer's disease-like *C. elegans* models. *STAR Protocols* **4** (2023).
17. Mahoney, T.R., Luo, S. & Nonet, M.L. Analysis of synaptic transmission in *Caenorhabditis elegans* using an aldicarb-sensitivity assay. *Nature Protocols* **1**, 1772-1777 (2006).
18. Davis, A.N. & Tanis, J.E. in *Journal of visualized experiments : JoVE* (2022).

#### Data availability

The mass spectrometry data reported in this paper is deposited to the ProteomeXchange Consortium via the PRIDE repository under accession number [accession number]. Any additional information required to re-analyze the data reported in this paper is available from the corresponding authors upon request.

#### Author contributions

Y.L., C.P., K.P. and L.J.R. conceived the study. Y.L., C.P., C.V., K.P. and L.J.R. designed the experiments. L.J.R. and K.P. supervised the study. Y.L., C.P., K.P. and L.J.R. wrote the manuscript. All authors reviewed and approved the final version of the manuscript.

Y.L. performed neuronal-related experiments and analyzed the mass spectrometry and other experimental data. C.P., K.K. and N.K. performed the *C. elegans* experiments and analyzed the resulting data. C.V. performed the primary mouse neuron experiments and related data analyses. W.T. and S.G. generated the *BNIP3L*-knockout iPSC lines. C.C. assisted with the Seahorse experiments. Z.Q.L. assisted with the confocal microscopy experiments. N.C. and L.C. assisted with data analysis. Z.Y. and A.J.N. performed sample preparation for mass spectrometry and analyzed the raw mass spectrometry data. V.A.B. assisted with establishing the iPSC cultures and contributed to project discussions. S.C. and Z.C. provided the iPSC lines. K.F. provided the iPSC lines and contributed to project discussions.

#### Funding statement

This work was supported by research grants from Nordea-fonden (L.J.R.), Novo Nordisk Foundation Challenge Programme: Harnessing the Power of Big Data to Address the Societal Challenge of Aging NNF17OC0027812 (L.J.R.), Aase og Ejnar Danielsens Fond (Y.L.), L.J.R. is a member of the Clinical Academic Group: Recovery Capacity After Acute Illness in Aging Population (RECAP). KP is funded by grants from the European Research Council (no. ERC-101077374 SynaptoMitophagy), the Fondation Santé and the Hellenic Foundation for Research and Innovation (HFRI; 23555).

#### Acknowledgements

We thank the Core Facility for Integrated Microscopy and the Flow Cytometry & Single Cell Core Facility at the Health and Medical Faculty at the University of Copenhagen for technical support. We thank Taylah Lea and Camilla Charlotte Nielsen Scheele from CBMR at University of Copenhagen for their kind help with Maestro MEA platform (Axion BioSystems).

#### Competing interests

The authors declare no competing interests.
